# N^α^-terminal acetylation of proteins by NatA and NatB serves distinct physiological roles in *Saccharomyces cerevisiae*

**DOI:** 10.1101/843953

**Authors:** Ulrike A. Friedrich, Mostafa Zedan, Bernd Hessling, Kai Fenzl, Ludovic Gillet, Joseph Barry, Michael Knop, Günter Kramer, Bernd Bukau

## Abstract

N-terminal (Nt)-acetylation is a highly prevalent co-translational protein modification in eukaryotes, catalyzed by at least five Nt-acetyltransferases (Nat) with differing specificities. Nt-acetylation has been implicated in protein quality control but its broad biological significance remains elusive. We investigated the roles of the two major Nats of *S. cerevisiae*, NatA and NatB, by performing transcriptome, translatome and proteome profiling of *natA*Δ and *natB*Δ mutants. Our results do not support a general role of Nt-acetylation in protein degradation but reveal an unexpected range of Nat-specific phenotypes. NatA is implicated in systemic adaptation control, as *natA*Δ mutants display altered expression of transposons, sub-telomeric genes, pheromone response genes and nuclear genes encoding mitochondrial ribosomal proteins. NatB predominantly affects protein folding, as *natB*Δ mutants accumulate protein aggregates, induce stress responses and display reduced fitness in absence of the ribosome-associated chaperone Ssb. These phenotypic differences indicate that controlling Nat activities may serve to elicit distinct cellular responses.

## Introduction

Acetylation of the N-terminal α-amino group of polypeptides (Nt-acetylation) is the most prevalent irreversible modification of proteins in eukaryotes, affecting 50–70% of the yeast proteome and up to 90% in higher eukaryotes (van Damme et al., 2011b; Helbig et al., 2010; Aksnes et al., 2016; Aksnes et al., 2019). Nt-acetylation is achieved already early during protein synthesis, as soon as the nascent chains emerge at the ribosomal tunnel exit (Gautschi et al., 2003), and therefore is an intrinsic property of proteins synthesized in eukaryotic cells. Co-translational Nt-acetylation is exerted by a set of structurally related N-terminal acetyltransferases (Nat) which associate with the large ribosomal subunit close to the peptide tunnel exit (Polevoda et al., 2008; Gautschi et al., 2003). *Saccharomyces cerevisiae* expresses five N-terminal acetyltransferases (NatA to NatE) with distinct specificities for N-termini (Aksnes et al., 2016), whereby NatA and NatB together are responsible for about 90% of all protein Nt-acetylation. NatA acetylates proteins with S-, A-, V-, G-, T- and C-termini, once the N-terminal methionine has been removed by methionine aminopeptidase (MetAP; Polevoda & Sherman, 2003b). NatB acetylates the N-terminal methionine of proteins starting with MD-, ME-, MQ-, and MN-(Polevoda & Sherman, 2003b). The extent of Nt-acetylation is variable for different proteins, ranging from infrequent acetylation to full acetylation, in particular for NatB substrates (Aksnes et al., 2016). NatA and NatB form each a dimeric complex consisting of a catalytic subunit and an auxiliary subunit that mediates ribosome binding (Polevoda et al., 2008; Gautschi et al., 2003). In yeast, the ablation of either subunit of NatA and NatB completely abolishes the enzymatic activity of the complex (Park & Szostak, 1992; Polevoda et al., 2003; Polevoda & Sherman, 2003a) and causes a number of Nat-specific and general phenotypes, including the NatA-specific mating defects, and a general reduction of growth rate as well as increased stress sensitivity, especially against elevated temperatures (Gautschi et al., 2003; Polevoda et al., 2003; Polevoda et al., 1999; Mullen et al., 1989; Whiteway et al., 1987). In searches for functional roles of N-acetylation, previous studies have implicated Nt-acetylation in a range of different cellular processes (Aksnes et al., 2019). First, Nt-acetylation is required for establishing protein-protein interactions, by providing critical contributions to involved interfaces. Examples are the Ubc12-Dcn1 ubiquitin ligase complex (Scott et al., 2011) and interactions of the silencers Sir3/ Orc1 with histone proteins (Whiteway et al., 1987; Geissenhöner et al., 2004; Wang et al., 2004), of the GTPase Arl3 with the Golgi membrane protein Sys1 (Behnia et al., 2004; Setty et al., 2004) and of actin with tropomyosin (Polevoda et al., 2003; Singer & Shaw, 2003). Second, Nt-acetylation contributes to the correct cellular sorting of proteins to the secretory pathway, by preventing the incorrect targeting of cytosolic proteins to the ER translocon (Forte et al., 2011). Third, Nt-acetylation has been connected to protein folding and stability since it reduces the N-terminal charge and thereby can stabilize N-terminal α-helices of proteins such as mitochondrial chaperonin 10 (Cpn10; Jarvis et al., 1995; Ryan et al., 1995), tropomyosin (Greenfield et al., 1994) and α-synuclein (Dikiy & Eliezer, 2014; Bartels et al., 2014). Finally, a number of publications link Nt-acetylation to protein degradation in yeast (Oh et al., 2017; Dörfel et al., 2017; Holmes et al., 2014; Zattas et al., 2013; Pezza et al., 2009). There, acetylated N-termini form an N-degron that targets proteins via the Ac/N-end rule pathway for ubiquitination by E3 ubiquitin ligases including Not4 and Doa10, followed by proteasomal degradation (Hwang et al., 2010; Shemorry et al., 2013). It has been proposed that the failure to form protein complexes exposes acetylated N-termini of unassembled subunits for ubiquitination via the Ac/N-end rule pathway, defining Nt-acetylation as mark for cellular control of protein stoichiometries and assembly. In support of this proposal, several different proteins overproduced from plasmids undergo degradation via the Ac/N-end rule pathway under specific conditions (Hwang et al., 2010; Shemorry et al., 2013). However, a recent study systematically analyzing the impact of Nt-acetylation on the stability of proteins in yeast did not provide evidence for a general function as N-degron (Gawron et al., 2016; Kats et al., 2018). The importance of Nt-acetylation for protein degradation *in vivo* is therefore still elusive.

To elucidate the biological functions of Nt-acetylation of proteins we performed a comprehensive multi-level analysis of mutant yeast cells lacking NatA or NatB. Combining transcriptome analysis, ribosome profiling, SILAC- and SWATH-based quantitation of protein stability and aggregation, we find that deletions of NatA and NatB cause distinct phenotypes that do not correlate with, and hence are not explained by quantitative contributions of NatA and NatB to Nt-acetylation of proteins. Our data suggest an unexpected divergence between the two Nat enzymes in the maintenance of cellular physiology. NatA activity is implicated in genetic control of Sir3 and Orc1 mediated gene silencing, transposon activity and mitochondrial activity. NatB activity rather supports protein homeostasis more globally, without major effects on protein stability but potentially by supporting the folding of newly synthesized proteins.

## Results

### Lack of NatA and NatB activity leads to distinct changes of gene expression profiles

We determined whether the absence of NatA or NatB affects gene expression and protein levels by a series of multi-omics high-throughput screens, comparing yeast wild-type (WT) with mutant cells lacking the catalytic subunits Naa10 or Naa20 of the NatA and NatB complexes (referred to as *natA*Δ and *natB*Δ), respectively. mRNA sequencing (RNAseq), ribosome profiling (RP) and quantitative proteomics allowed quantitative assessment of Nat-dependent changes at the transcriptional, translational and protein steady-state level. The deep sequencing-based data sets (RNAseq and RP) comparing *natA*Δ or *natB*Δ and WT include more than 4700 quantified genes and are highly reproducible among replicates (r = 0.79-0.95; **Figures S1A-S1D and S1F-S1H**). Stable isotope labeling by amino acids in cell culture (SILAC; Ong et al., 2002) coupled with mass spectrometry allowed to determine the steady-state levels of 2169 and 2976 proteins in *natA*Δ and *natB*Δ mutant cells, respectively (r = 0.68/0.71; **Figures S1E and S1I**). To complement SILAC-MS analysis, we also employed label-free SWATH-MS (Gillet et al., 2012) and determined the steady-state levels of 2245 (*natA*Δ) and 2220 (*natB*Δ) proteins.

For 78 proteins, among them 18 previously unknown Nt-acetylated proteins, we verified the expected, Nat-specific loss of Nt-acetylation in the *natA*Δ and *natB*Δ mutants (**Figures S2A and S2B**). Our data confirm the lack of functional redundancy between NatA and NatB. Supporting previous findings (van Damme et al., 2011a; Aksnes et al., 2016), NatB substrates with MD-, ME-, MN- and MQ-termini are acetylated at efficiencies close to 100%, while some of the A-, G-, S- and T-termini of NatA substrates are only partially Nt-acetylated.

Loss of Nt-acetylation in *natA*Δ and *natB*Δ mutants causes substantial, mutant-specific changes in protein synthesis. RP analysis revealed that the deletion of NatA and NatB reproducibly changes the translation levels of approximately 150 and 400 genes more than two-fold, respectively (**Figures 1A and 1B**). The changes are only weakly correlated between the two mutants (r = 0.182; **Figures S3B and 1C**). Consistent with the more severe growth defects of *natB*Δ mutant cells on agar plates (**Figure S4**), the extent of changes is overall stronger in *natB*Δ cells as compared to *natA*Δ cells.

**Figure 1.**
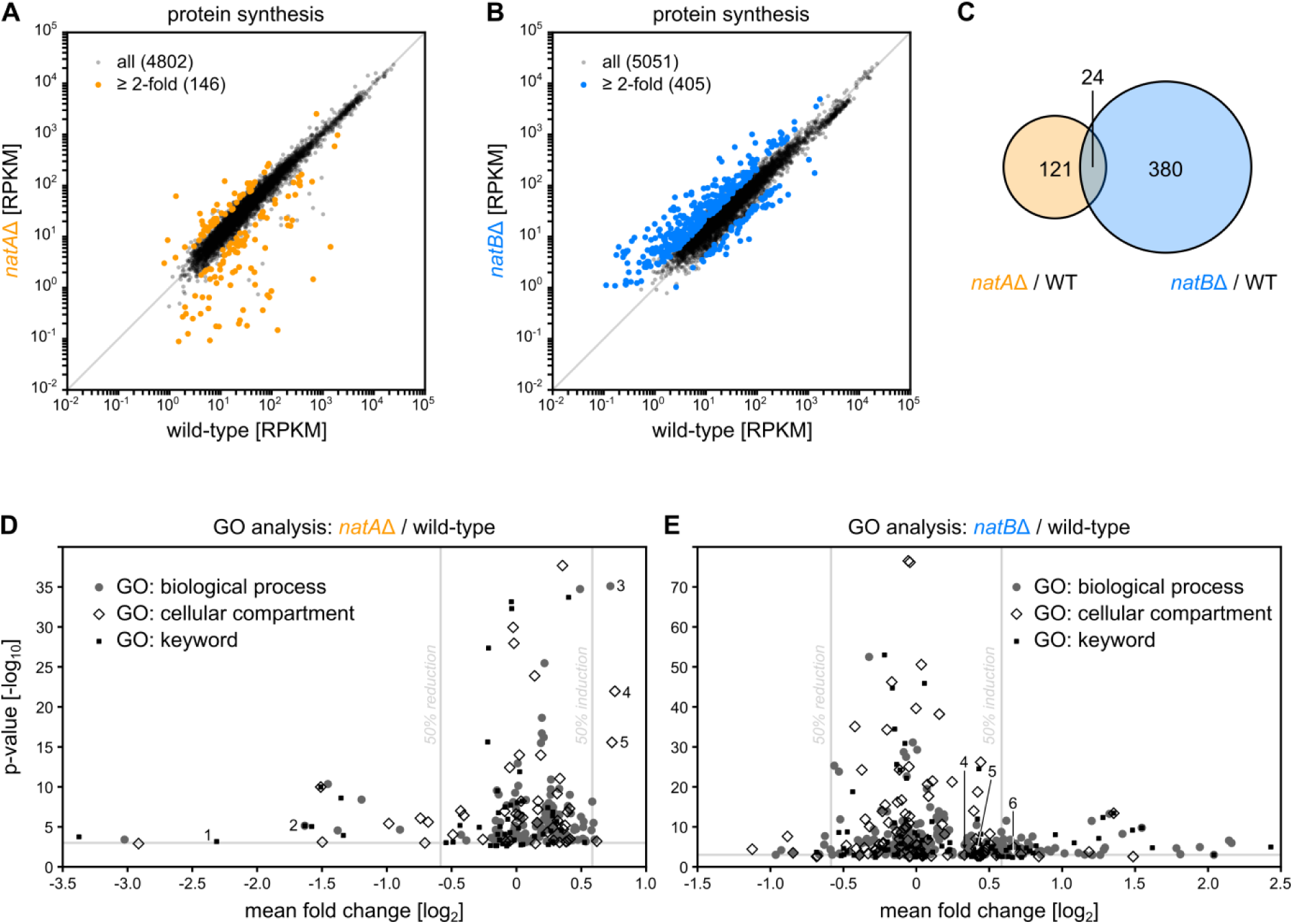
Changes in translation upon deletion of NatA or NatB. Levels of translation of open reading frames are compared between WT and *natA*Δ (A) or *natB*Δ (B) cells using ribosome profiling. Replicate 1 is shown; n = 2. (C) Comparison of differentially expressed genes between *natA*Δ and *natB*Δ mutant cells. (D-E) Threshold-independent gene ontology (GO) analysis based on results from (A) and (B), respectively. (FDR = 0.02) Indicated GO terms: 1 – Pheromoneresponse, 2 – viral procapsid maturation/ viral reproductive process/ Capsidmaturation/ others, 3 – mitochondrial organization, 4 – mitochondrial/ organellar large ribosomal subunit, 5 – mitochondrial/ organellar small ribosomal subunit, 6 – Stressresponse.

The changes in protein synthesis in *natA*Δ or *natB*Δ strongly correlate with changes in mRNA levels as revealed by RNAseq analysis (r = 0.94/0.87; **Figures S5A and S5D**). This demonstrates that they predominately result from changes at the transcriptional level. We also found good correlations between observed changes in protein synthesis rates and steady-state levels determined by SILAC (r = 0.64/0.49; **Figures S5B and S5E**). The observed changes of the proteome are therefore mainly due to altered protein synthesis rather than changed protein degradation.

We performed threshold-independent gene ontology (GO) enrichment analyses to identify the gene expression responses to deficiencies in Nt-acetylation activity in the respective mutants (**Figures 1D and 1E**). Most prominently, the lack of NatA activity impacts the expression of pheromone response genes, transposable elements and genes encoding mitochondrial ribosomal proteins (**Table S1**). The lack of NatB activity particularly impacts the expression of genes involved in stress-responses, amino acid biosynthesis and translation (**Table S2**).

### NatA regulates gene silencing and mitochondrial protein synthesis

The most strongly down-regulated genes in *natA*Δ cells are enriched for genes encoding mating type-related proteins, especially MATa-specific proteins (**Figures 2A and 2B**). Haploid yeast cells sense mating pheromone as signal for the presence of potential mating partners. This triggers the mating pathway to induce cell cycle arrest at G_1_ phase and oriented growth towards the mating partner for cell fusion (Bardwell, 2004; Haber, 2012). Some members of this signaling cascade are expressed in both yeast mating types, MATa and MATα, while others are mating type-specific. In diploid cells, all mating genes are suppressed by the combined, dimeric repressor Mata1-Matα2 derived from both mating type loci, *HML* (MATα) and *HMR* (MATa).

**Figure 2.**
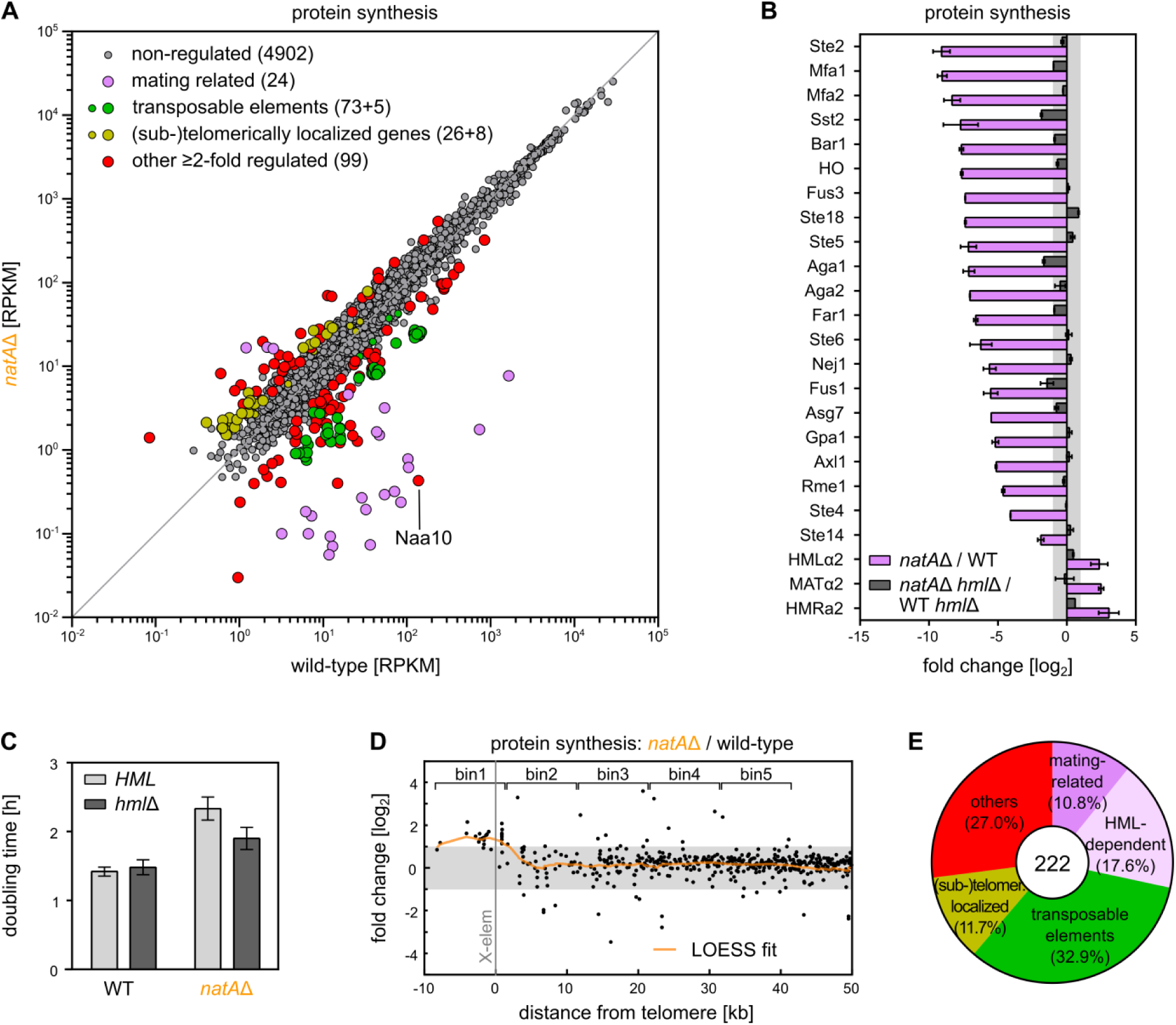
NatA-dependent regulation of gene silencing at the *HML* locus and near telomeres. (A) Protein synthesis is compared between WT and *natA*Δ mutant cells grown in YPD media using ribosome profiling. Replicate 1 is shown; n = 2. Significantly enriched or disenriched protein groups are colored. Naa10 represents the catalytic subunit of NatA deleted in the *natA*Δ strain. (B) Altered protein synthesis of MATa-specific proteins, as labeled in (A), is shown comparing the *HML*^+^ and *hml*Δ background. (C) Growth analysis of *natA*Δ and *hml*Δ mutant cells in YPD media. (D) The fold-change of synthesis of each protein determined by ribosome profiling is displayed in respect to its gene localization on chromosomes. (E) Quantification of ≥ 2-fold differentially synthesized proteins corresponding to their function/ gene localization.

We found that MATa-specific genes are almost completely repressed upon NatA deletion, providing a rationale for the strongly impaired mating capacity of MATa *natA*Δ cells (Whiteway & Szostak, 1985). We hypothesized that deficient Nt-acetylation causes these alterations in gene expression through effects on the gene silencer Sir3, which confers suppression of gene expression at the silent *HML* locus of the mating cassette (Rine & Herskowitz, 1987). Nt-acetylation of Sir3 is required for its stable interaction with the nucleosome core particle during gene silencing (Yang et al., 2013). Confirming this assumption, deletion of the *HML* cassette rescued expression of MATa-specific genes in *natA*Δ cells (**Figures 2B and S6A**). Intriguingly, it also restored expression of all five families of transposable elements which are downregulated in *natA*Δ mutant cells (**Figures 2A and S6A**). Considering that transposable elements are exclusively expressed in haploidic cells (Elder et al., 1981), we speculate they may be regulated similar to the *MATa*-specific mating response. Our gene expression analysis in *natA*Δ cells identified 41 additional, so far unidentified *HML*-dependent genes with unrelated functions (**Table S3**).

Of note, the additional *HML* deletion rescued expression of more than half of the differentially expressed genes in *natA*Δ mutant cells, i.e. transposable elements, MATa-specific genes and others (**Figure S6A**), and this also partially alleviates the NatA-dependent slow growth phenotype (**Figures 2C and S6B**).

An additional set of NatA-regulated genes is differentially expressed independently of the *HML* locus and predominantly localized close to sub-telomeric regions (**Figures 2D, 2E, S6D, and S6E**). Intriguingly, sub-telomeric genes are typically repressed by the silencer Orc1, a NatA substrate that is structurally homologous to Sir3, possesses an identical N-terminus and also functions similarly (Geissenhöner et al., 2004). The de-repression of sub-telomeric genes is specific for *natA*Δ mutants in comparison to mutants affecting other N-terminal acetyltransferases (**Figure S6C** and data not shown). These findings strongly suggest that Nt-acetylation of Orc1 is critical for its gene silencing activity.

Based on our RNAseq, RP and SILAC-MS analyses, NatA deletion also elevated the average synthesis of nuclear-encoded mitochondrial ribosomal proteins by more than 40% (**Figures 3A and S7A**). This effect was much more pronounced in *natA*Δ mutants as compared to *natB*Δ mutants, suggesting specificity of NatA versus NatB activity (**Figure S7B**). 94% of all detected genes encoding mitochondrial ribosomal proteins are up-regulated in *natA*Δ cells (76/81 in the RP data set), leading to similarly elevated ribosomal protein levels (**Figure 3A**). The changes in both, ribosomal protein expression and mitochondrial phenotype were not affected by mating-type regulation of the *HML* cassette (**Figures S7A, S7C, and S7D**).

**Figure 3.**
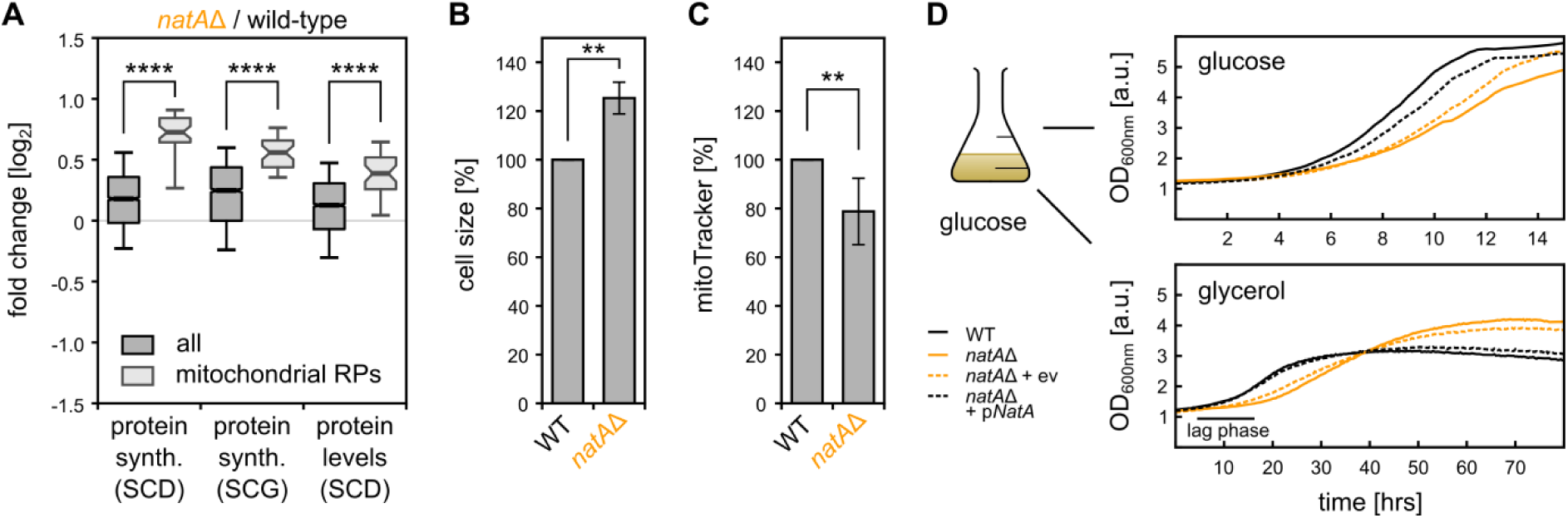
Effect of NatA on mitochondrial ribosome levels and mitochondrial activity. (A) Synthesis and steady state levels of proteins in cells grown in synthetically defined media containing glucose (SCD) and glycerol (SCG) as carbon source determined by ribosome profiling and SILAC, respectively. Log_2_-transforms of all proteins (2372) and mitochondrial ribosomal proteins (38) are statistically compared by Mann-Whitney U test. FACS-based analysis of (B) cell size and (C) mitoTracker intensity, normalized to the cell size. 6 biological replicates are combined and compared using Mann-Whitney U test. (D) Growth curves of WT and *natA*Δ mutant cells as indicated. Cells of the indicated strains were grown in glucose-based SC media to mid-log phase and then shifted to different media (start OD = 0.1) after careful washing. ev – empty vector.

The seemingly elevated protein synthesis capacity in mitochondria however did not coincide with enhanced respiration capacity of *natA*Δ cells analyzed by MitoTracker. Correcting the measured respiration activity for the larger cell size of NatA-depleted cells even suggests reduced mitochondrial activity (**Figures 3B and 3C**). In line with this finding, *natA*Δ cells displayed a greatly extended lag phase when shifted from the fermentable carbon source glucose to respiration-obligate glycerol (**Figure 3D**). Together our data suggest that enhanced synthesis of mitochondrial ribosomal proteins may be induced in response to impaired function of mitochondria or mitochondrial translation in *natA*Δ cells.

### Constitutive stress response in *natB*Δ mutant cells

Our differential gene expression analysis revealed that *natB*Δ cells strongly up-regulate genes encoding machinery for protein refolding, including the stress-inducible chaperones Hsp26 and Hsp70 (Ssa4) as well as the cytosolic and mitochondrial disaggregases Hsp104 and Hsp78 (**Figure 4A**). These genes were only poorly induced in *natA*Δ cells, indicating that the lack of NatB activity, but not of NatA activity, causes significant protein folding stress eliciting cell responses (**Figure 4B**). Consistent with the elevated expression of chaperone genes, *natB*Δ cells more efficiently refolded heat-denatured luciferase after thermal stress treatment of the cells as compared to *natA*Δ cells (**Figure 4C**). Interestingly, genes more than two-fold induced in *natB*Δ cells were enriched for targets of the transcription factors Msn2/Msn4 which control expression of the general stress response of yeast (Martínez-Pastor et al., 1996; Schmitt & McEntee, 1996) (**Figure 4D**). Furthermore, in *natB*Δ cells the burden of unfolded proteins on the cellular protein folding machinery is counteracted by reduced synthesis of proteins involved in cytosolic translation (**Figure 4E**). The reduction of translation activity was also observed by polysome profiling that showed a higher monosome to polysome ratio in *natB*Δ cells as compared to WT and *natA*Δ cells (**Figure 4F**). In agreement with the reduction in growth rate, ribosome synthesis and translation activity, *natB*Δ mutant cells also showed an about 2-fold lower rate of incorporation of radiolabeled methionine into newly synthesized proteins as compared to WT cells (**Figure 4G**).

**Figure 4.**
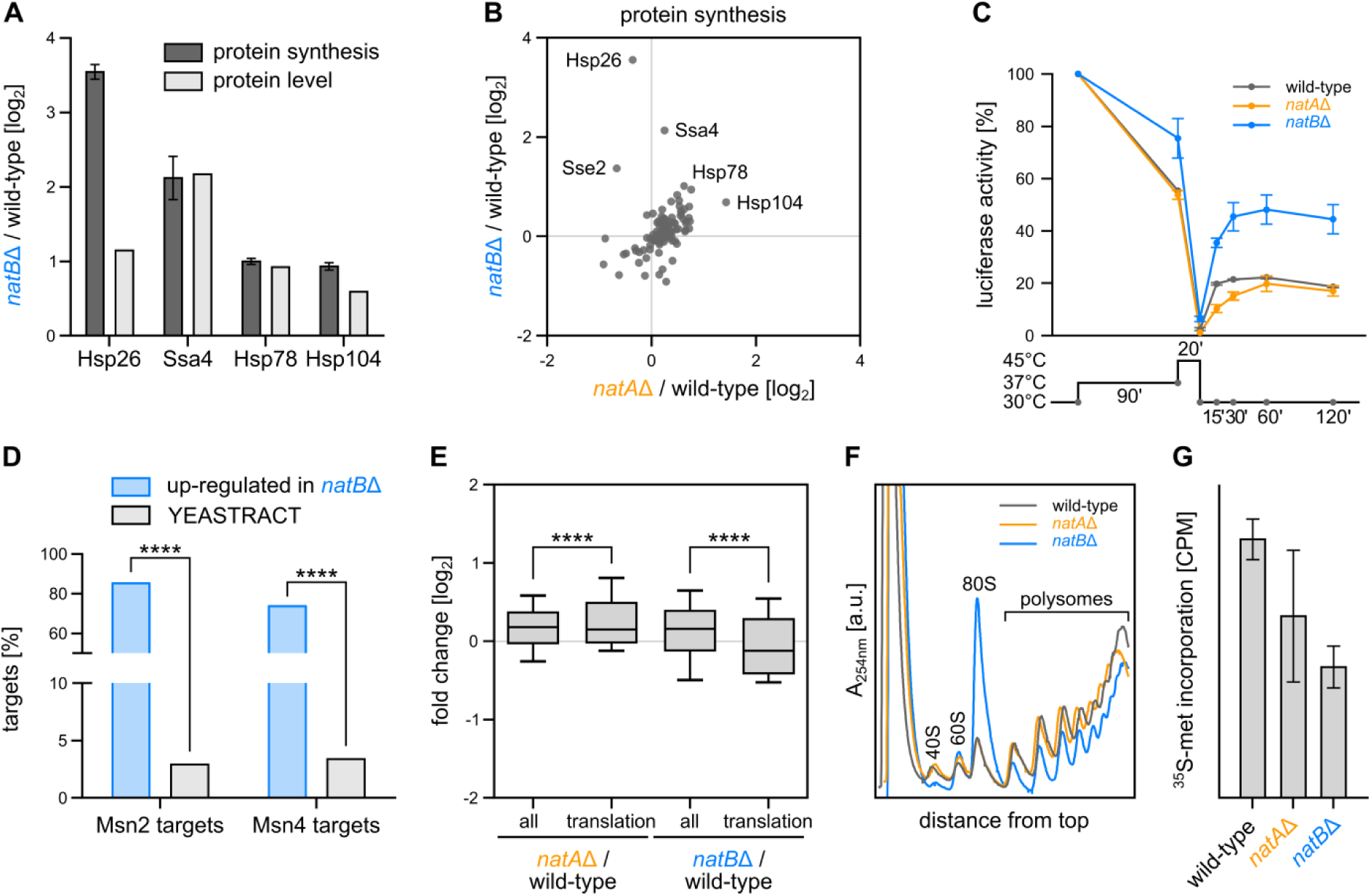
Constitutively induced stress response in *natB*Δ cells. (A) Changes in *natB*Δ and WT cells of synthesis (RP) and steady state levels (SWATH proteomics) of chaperones involved in protein refolding. (B) Changes of synthesis (RP) of the protein folding machinery (GO: protein folding) in *natA*Δ and *natB*Δ mutant cells relative to WT. (C) *In vivo* refolding kinetics of heat denatured luciferase expressed in WT, *natA*Δ and *natB*Δ mutant cells. The heat shock regime used is indicated above. Cycloheximide was added prior to the heat shock. (D) Transcription factor analysis of upregulated genes (>2-fold) in *natBΔ* cells using the “rank by transcription factor” algorithm of the YEASTRACT database that compares the percentage of transcription factor targets in the dataset and the YEASTRACT database. Msn2 and Msn4 are the two top ranked transcription factors associated with the upregulated genes. (E) Comparison of the distribution of protein synthesis levels (RP) for proteins annotated to “translation” for *natA*Δ and *natB*Δ mutant cells relative to WT. (F) Polysome profiles of *natA*Δ and *natB*Δ mutant and WT cells performed by fractionation of cell lysates in 10-50% sucrose gradients. (G) ^35^S-methionine incorporation after 10 min pulse labeling in TCA-precipitated proteins in *natA*Δ and *natB*Δ mutant and WT cells (measured in counts per million (CPM) normalized to OD_600nm_).

### Lack of Nt-acetylation has no global impact on substrate stability

The exposure of acetylated N-termini was recently described as the targeting signal for degradation of surplus or unassembled protein complex subunits via the Ac/N-end rule pathway (Hwang et al., 2010; Shemorry et al., 2013). Accordingly, the constitutively upregulated stress response observed in *natB*Δ cells, and to a smaller extent in *natA*Δ cells, may be triggered by the accumulation of non-acetylated Nat substrates that fail to be degraded. However, a series of independent experiments do not support this idea. First, the average steady-state levels of both, verified NatA and NatB substrates and all cellular proteins did not differ when comparing WT and *natA*Δ or *natB*Δ cells grown at physiological conditions at 30°C (**Figures 5A and 5B**, first panels). Second, as revealed by two independent approaches, protein stabilities are similar for Nat substrates versus all proteins. In one approach, we calculated protein stabilities by normalizing protein levels (determined by quantitative proteomics) to synthesis rates (determined by RP). The results showed that global protein stabilities are not affected by deletion of NatA or NatB (**Figures 5A and 5B**, second panels). In a second approach, focusing on *natB*Δ mutant cells, we measured protein stabilities using the recently developed tandem fluorescent protein timer approach (tFT; Khmelinskii et al., 2012) and compared *natB*Δ mutant to WT cells. The tFT comprises two single-color fluorescent proteins (sfGFP and mCherry) that mature with different kinetics and are fused to a protein of interest. Determining the ratio of mCherry to sfGFP allows measuring the relative protein stability, independent of protein abundance. Analyzing a genome wide library of chromosomally encoded tFT fusion genes did not show any significant difference of tFT-tagged verified NatB substrates as compared to all measured proteins, confirming previous results (**Figure 5C**, first panel). Third, the Ac/N-end rule pathway does not play a detectable role as quality control mechanism when protein homeostasis is perturbed by environmental stress. RP, SILAC and tFT analyses of cells transiently exposed to heat stress (at least two doublings at 37°C) did not detect global effects of NatA and NatB deletion on protein levels or stability (**Figures 5A-5C**, right panels). Fourth, we did not detect a role of Nt-acetylation on protein subunit degradation when subunit stoichiometries of specific protein complexes are disturbed. This was experimentally investigated by duplicating genes encoding individual subunits of complexes, fusing one of the alleles to the tFT-encoding sequence, and measuring the stability of the tagged subunits. The 2-fold molar excess of the subunits encoded by the gene duplications over the other subunits of the corresponding protein complex should lead to an experimentally detectable degradation of up to 50% of the tagged subunits. The chosen candidates are Nt-acetylated members of oligomeric complexes, with two of them (Hsp104 and Ubp6) being positively tested in a previous study for stability changes under strong overexpression (Hwang et al., 2010). Deficient Nt-acetylation in *natA*Δ and *natB*Δ cells did not increase the stability of the excess subunits. Ubp6 and Sup45 were destabilized in both mutant cells, indicating that stability was independent of Nt-acetylation. The only exception is Hsp104, which is slightly stabilized in *natB*Δ mutants (**Figure 5D**).

**Figure 5.**
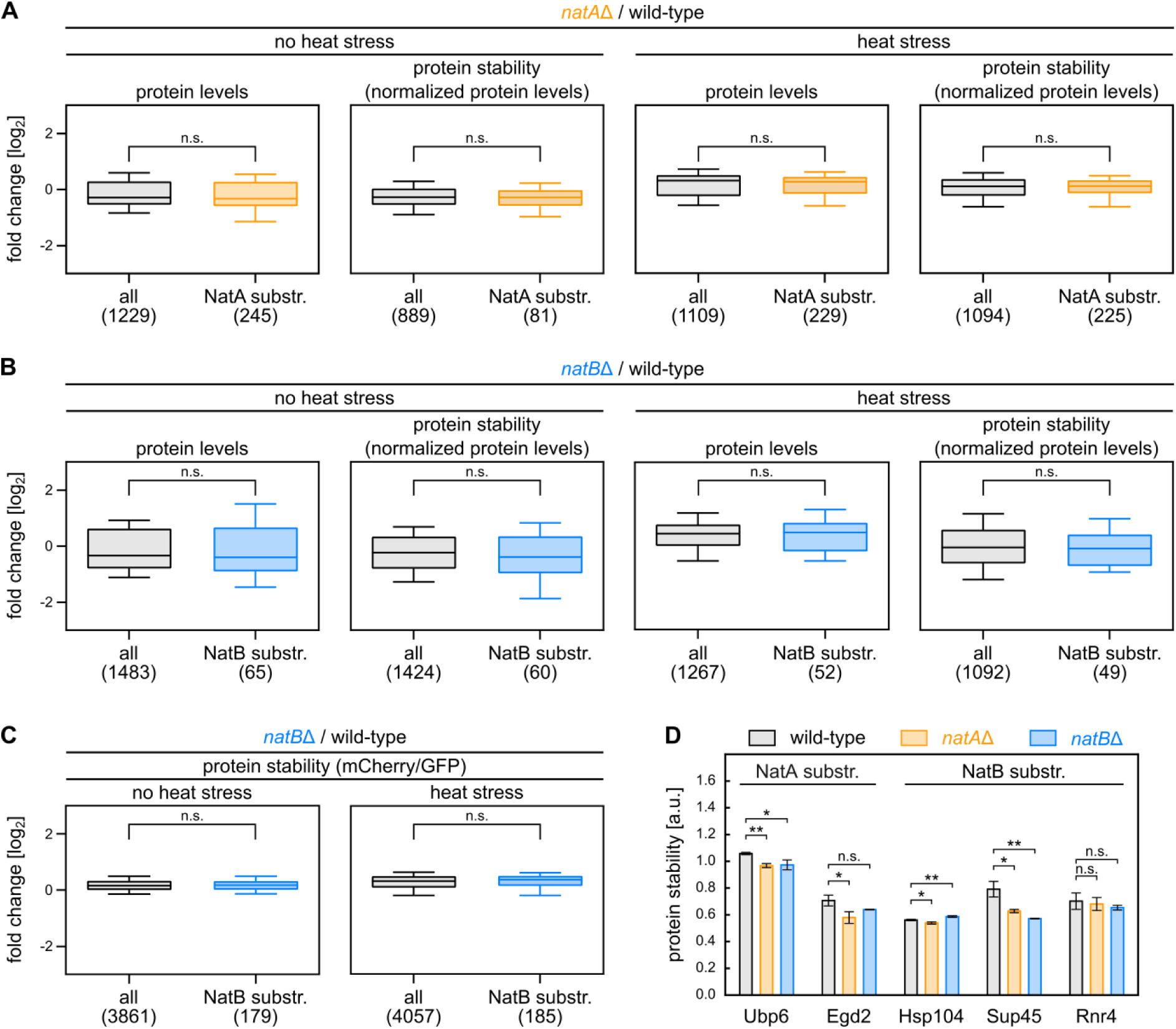
Loss of Nt-acetylation has no global impact on substrate stability. Protein levels of *natA*Δ (A) and *natB*Δ (B) mutant relative to WT cells based on quantitative proteomics (SILAC) and protein stabilities as normalized protein levels (SILAC relative to protein synthesis determined by RP). (C) Stability of tFT-tagged proteins in *natB*Δ mutant relative to WT cells measured using the mCherry/GFP ratio as read-out. Cells were grown under physiological (30°C) and heat stress (37°C for two doubling times (SILAC) or 1 day (tFT)) conditions. Verified Nt-acetylation substrates are compared versus all quantified proteins. The number of proteins in each group is indicated between brackets. (D) Comparison of the stability of tFT-tagged proteins (mCherry/GFP as read-out) expressed in WT and *natA*Δ and *natBΔ* mutant cells.

Taken together, our analyses do not indicate a prominent role of Nt-acetylation for global protein stability at physiological and heat stress conditions and upon perturbance of protein stoichiometries as occurring upon gene duplication.

### Global protein aggregation upon lack of Nt-acetylation

Another possible reason for the induction of the protein refolding machinery in *natB*Δ cells (**Figure 4A**) would be that the loss of Nt-acetylation compromises the structural integrity of NatB substrates, necessitating the refolding and sequestration activity of chaperones and ultimately resulting in increased protein aggregation. To address this possibility, we isolated protein aggregates from *natA*Δ and *natB*Δ mutant and WT cells grown at 30°C. Absence of NatB, and to a smaller extent also of NatA, led to enhanced aggregation of a diverse set of endogenous proteins (**Figure 6A**). Employing SILAC-based quantitative proteomics analysis of the fraction of aggregated proteins, we identified 613 proteins that are more than 2-fold enriched in the aggregate fraction in *natA*Δ relative to WT, and 794 proteins in *natB*Δ cells (**Figure 6B**). The majority of the aggregating proteins in the mutants had no or only modest changes in their total levels in comparison to WT, indicating that protein aggregation is not driven by altered total protein levels in the mutants (**Figures S8A and S8B**). Unexpectedly, the two sets of aggregated proteins in *natA*Δ and *natB*Δ cells largely overlap (54-70%; **Figure 6B**). This implies a general folding deficiency conferred by deficiency in Nt-acetylation that is not specific to NatA or NatB substrates. Consistently, protein aggregates in *natA*Δ and *natB*Δ cells were not enriched for the specific substrates of the missing Nat enzyme (**Figure S9A**), and in the *nat*Δ mutant cells the aggregation propensity of all cellular proteins was not significantly different from that of Nat substrates (**Figure S9B**).

**Figure 6.**
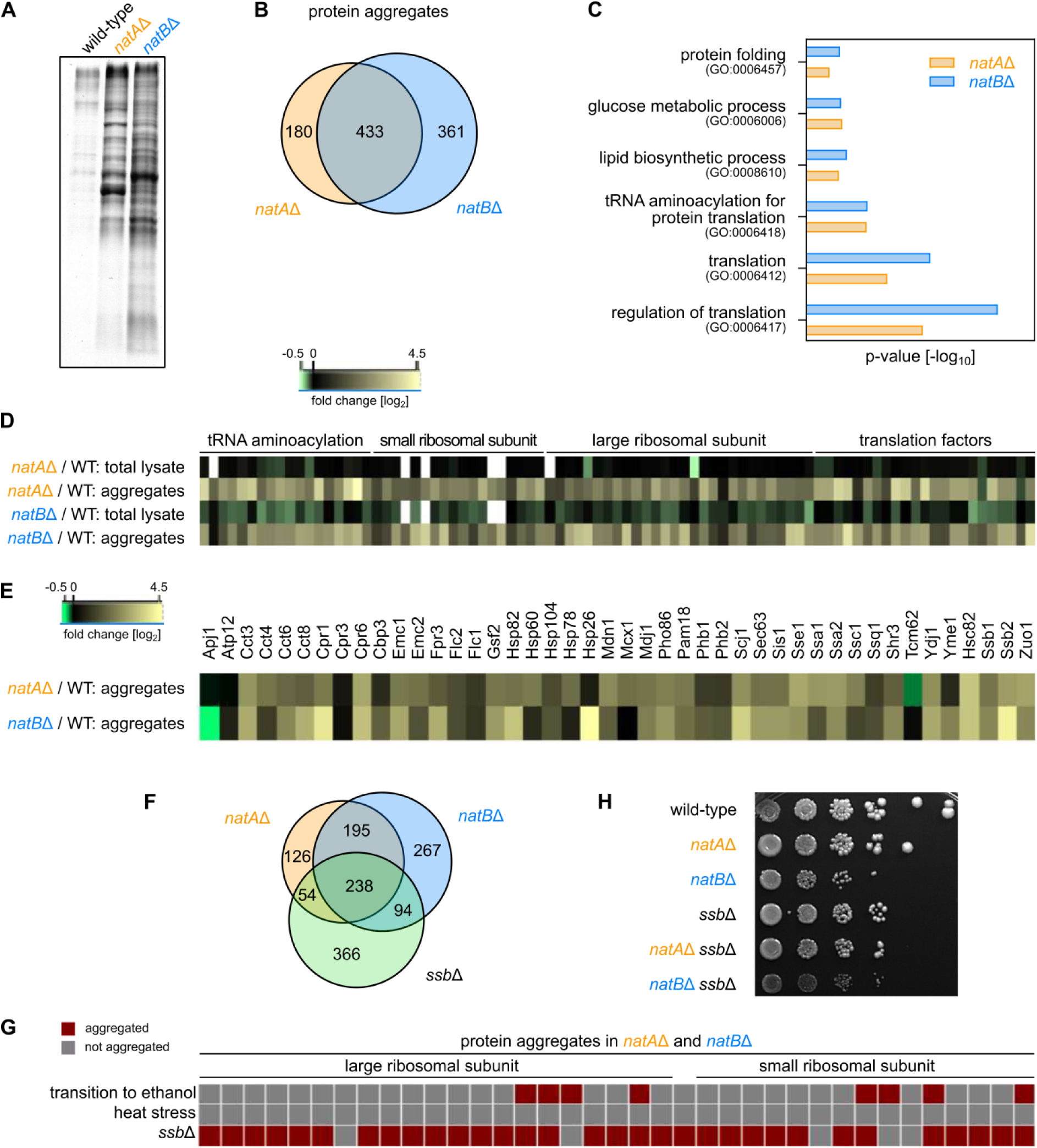
Global protein aggregation upon loss of N-terminal acetylation. (A) Isolated protein aggregates from WT, *natA*Δ and *natB*Δ mutant cells visualized by SDS-PAGE and Coomassie staining. (B) The overlap between protein aggregates in *natA*Δ and *natB*Δ mutant cells. (C) GO term enrichment analysis for biological processes of aggregated proteins in *natA*Δ *and natB*Δ mutant cells. Top GO terms based on p-value are shown. (D) Protein level changes between *natA*Δ, *natB*Δ and WT cells in the total lysate, and aggregate fraction for components of the cytoplasmic translation machinery. (E) Enrichment of proteins that mediate protein folding (GO: protein folding) in aggregates in *natA*Δ *and natB*Δ mutant cells, relative to WT. (F) The overlap between protein aggregates in *natA*Δ, *natB*Δ and *ssb*Δ mutant cells. (G) The overlap of aggregated ribosomal proteins in *natA*Δ and *natB*Δ mutant cells with aggregated proteins in *ssb*Δ mutant cells and WT cells subjected to different stress conditions (heat stress, transition from glucose to ethanol). (H) Growth of *ssb*Δ, *natA*Δ, and *natB*Δ double deletion mutants in comparison to single deletion mutants and WT on YPD agar plates at 30°C.

We surmised that protein misfolding and aggregation could also be caused by the inactivation of chaperones, due to impaired Nt-acetylation of chaperones themselves or to elimination of a so far unknown chaperone activity of the Nat enzymes. Arguing against the first possibility, the large majority of chaperones are substrates of NatA (e.g. Hsp26, Ssa1/2, Ssb1/2, Ssz1, Sse1/2, Egd2, Hsc82, Hsp82). The sole relevant exception is Hsp104, which is a substrate of NatB. However, the chaperone activity of Hsp104 is restricted to the solubilization of aggregated proteins (Mogk et al., 2018), and this activity (as measured for aggregated luciferase) was even about 2-fold elevated in *natB*Δ cells (**Figure 4C**). These arguments strongly suggest that protein aggregation in *natB*Δ cells cannot be explained by chaperone inactivation. To investigate the hypothesis that Nat complexes may function as chaperones, we tested the capacity of a previously described catalytically inactive NatA-Naa10 mutant (Liszczak et al., 2013) to suppress protein aggregation in *natA*Δ cells. However, the expression of plasmid-encoded Naa10^E26A^ mutant protein did neither prevent protein aggregation nor revert the *natA*Δ growth phenotypes, in contrast to the expression of NatA WT as control (**Figures S10A and S10B**). This result argues against this possibility and suggests that the enzymatic activity of Nt-acetylation is critical for NatA functionality in protein homeostasis *in vivo*.

We next searched for enriched biological processes involving the fraction of aggregated proteins in *natA*Δ and *natB*Δ cells, by performing a threshold-dependent GO analysis. The most enriched group among the aggregated proteins in *natA*Δ and *natB*Δ mutants are cytoplasmic translation components including cytosolic small and large ribosomal subunits, translation factors, and tRNA-aminoacylation machinery (**Figures 6C and 6D**). Although a number of aggregating translation factors are known stress granule markers, we did not detect any stress granule formation in *nat*Δ mutant cells. This suggests that impaired Nt-acetylation reduces synthesis and selectively inactivates and sequesters components of the protein synthesis machinery, which further reduces the flux of newly synthesized proteins into the network of chaperones and factors involved in nascent polypeptide maturation. As expected, chaperones implicated in protein folding and disaggregation were also enriched in protein aggregates in *natA*Δ and *natB*Δ cell (**Figures 6C and 6E**).

Intriguingly, the aggregation-prone proteins that we identified in Nat-deficient mutants are highly similar to those isolated from cells lacking the ribosome associated Hsp70s, Ssb1 and Ssb2 (Koplin et al., 2010) (**Figure 6F**). This overlap includes many ribosomal proteins, which do not aggregate under general stress conditions (**Figure 6G**). Furthermore, deletion of *SSB1/2* in *natB*Δ cells strongly reduced cellular fitness (**Figure 6H**), suggesting Ssb1/2 and NatB act in parallel pathways. Given that Nat enzymes and Ssb1/2 both act co-translationally, we tested whether the loss of Nt-acetylation has a particularly strong impact on the *de novo* folding, and hence the structural integrity of nascent proteins, by radioactive pulse-labeling of newly synthesized proteins followed by aggregate isolation. Indeed, aggregates isolated from *natB*Δ cells contained elevated amounts of ^35^S**-**methionine labeled proteins as compared to WT, suggesting that non-acetylated nascent proteins have an increased tendency to misfold and aggregate (**Figure S11**).

## Discussion

Our study provides new insights into the functional importance of Nt-acetylation of proteins. Multi-level high-throughput analysis of gene expression and proteome homeostasis of *natA*Δ and *natB*Δ mutant cells revealed that the major N-terminal acetyltransferases NatA and NatB have distinct roles in physiology and proteostasis of *S. cerevisiae*. Lack of NatB activity causes much stronger perturbation of protein homeostasis than the lack of NatA activity, resulting in misfolding and aggregation of several hundred proteins and induction of stress-induced genes controlled by the Msn2/4 transcription factors. Lack of NatA activity instead affects primarily gene expression, genome integrity and metabolic regulation. The disparate phenotypes cannot be explained by differences in substrate pool sizes of the two enzymes since the substrate spectrum of NatB is considerably smaller than that of NatA (3-fold less protein species, 13-fold lower number of substrate molecules, less complete acetylation of a given protein species; Aksnes et al., 2016). Rather, they appear to rely at least in part on a set of enzyme-specific targets, such as transcription silencers Sir3 and Orc1 acetylated by NatA. Our data furthermore suggest that for both NatA and NatB targets, the Ac/N-end rule degradation pathway is a less prominent cellular response to misfolding stress than proposed earlier (Hwang et al., 2010; Shemorry et al., 2013).

One possible explanation for the prominent protein aggregation in *natB*Δ mutants is that the loss of Nt-acetylation impairs the folding or destabilizes the folded state of non-acetylated proteins as previously shown for some endogenous proteins, e.g. chaperonin 10 (Ryan et al., 1995; Jarvis et al., 1995) and tropomyosin (Greenfield et al., 1994). However, we find that Nat-specific substrates are not enriched in the aggregate fraction of the respective mutant cells. The aggregate fraction is also not enriched in subunits of protein complexes, suggesting that the lack of Nt-acetylation of complex subunits is also not a major reason for the spectrum of aggregating proteins. Nat substrates that misfold and aggregate in *nat*Δ mutants due to the lack of Nt-acetylation may instead promote the co-aggregation of other proteins, thereby causing global disbalance of the proteome.

A second plausible mechanism explaining the aggregation phenotype is that *natB*Δ cells have reduced chaperone activity. This may result from (i) lack of Nt-acetylation of the chaperones themselves, or because (ii) Nat complexes have chaperone-like activities or (iii) Nat complexes affect the activity of chaperones (Ssb and NAC) that act co-translationally in the vicinity of Nat complexes at the ribosomal tunnel exit. The first possibility is unlikely since the vast majority of chaperones is Nt-acetylated by NatA, and hence retains functionality in *natB*Δ mutants. We performed tests of the second possibility for NatA but did not obtain evidence for a chaperone function. The most compelling evidence points towards the third possibility. Deletion of *SSB1/2* in *natB*Δ cells strongly diminished cellular growth, implying a functional overlap between co-translational folding assisted by Ssb and Nt-acetylation by NatB. Consistent with this scenario, aggregates isolated from pulse-labelled *natB*Δ cells include newly synthesized proteins, indicating that a fraction of them misfolds and becomes sequestered into aggregates. A functional overlap was previously suggested also for NatA and Ssb, based on the partial rescue of heat sensitivity and other *natA*Δ phenotypes by *SSB1/2* overexpression (Gautschi et al., 2003). Importantly however, *natA*Δ mutants lacking *SSB1/2* did not display a reduced cellular fitness that is comparable to *natB*Δ cells lacking *SSB1/2*. We speculate that Nat enzymes coordinate ribosome association and function of Ssb, thereby coordinating co-translational protein folding and enzymatic processing. Consequently, the lack of Nt-acetylation may impose challenges for the co-translational folding of proteins, which in turn enhances the need for co-translational folding assistance, explaining the negative effect of simultaneous deletion of NatB and Ssb on growth. Evidences for the pivotal role of NatB in folding of nascent proteins are (i) the aggregation of ^35^S**-**methionine labeled proteins in *natB*Δ cells after a short pulse of ^35^S**-**methionine, (ii) the overlap between protein aggregates in *natB*Δ and *ssb1,2*Δ cells, (iii) the impaired fitness of *natB*Δ *ssb1,2*Δ double deletion mutants, and (iv) the co-translational action of NatB, like Ssb1/2, via association with the ribosomal exit tunnel. Misfolding and aggregation of nascent chains in *natBΔ* cells may drive co-aggregation of ribosomes and other misfolding-prone proteins through unspecific protein-protein interactions. This could account for the observed proteostasis disbalance including induction of the heat shock response and overall reduced protein synthesis.

Nt-acetylation was proposed to confer a quality control pathway for unassembled orphan subunits, by creating a degron, a signal for ubiquitination and proteasomal degradation (Shemorry et al., 2013). Accordingly, the lack of Nt-acetylation should lead to accumulation of unassembled subunits, eventually triggering protein aggregation and stress responses, consistent with what we observed in *natB*Δ and to a smaller extent in *natA*Δ cells. However, our analyses of the effects of Nt-acetylation on protein levels and stability using quantitative proteomics, RP and tFT approaches do not provide further support for this model. We do not detect a significant increase of steady-state levels or half-lives of experimentally verified Nat substrates upon loss of Nt-acetylation, neither under physiological growth conditions nor after perturbation of protein folding and complex assembly by heat stress and gene duplication. Our results do not exclude that N-acetyl mediated degradation may control the stability of specific proteins, or control protein stability under specific conditions such as strong overproduction of Nt-acetylated orphan subunits. We also do not exclude the existence of redundant degradation pathways that target Nat substrates for degradation in the absence of Nt-acetylation, as shown previously for a subset of unacetylated NatC substrates (Kim et al., 2014).

Although the number and diversity of proteins acetylated by NatA is much higher than that of NatB (Aksnes et al., 2016; van Damme et al., 2011a), protein aggregation is less prominent in *natA*Δ cells, indicating non-random distribution of aggregation-prone substrates among the two enzymes. The most prominent phenotypic changes related to NatA ablation instead affect gene expression, genome integrity and metabolic regulation. NatA is specifically involved in gene silencing to control sexual differentiation and mating, and expression of mitochondrial genes. *natA*Δ cells, due to their incapability to Nt-acetylate Sir3, fail to suppress the *HML* locus and artificially express not only the MATa-specific genes *MATa1* and *MATa2* from the *HMR* locus but also the MATα*-*specific genes *MAT*α1 and *MAT*α2 from the *HML* locus, resulting in a pseudodiploid, sterile phenotype. The functional impairment of Sir3 in *natA*Δ cells most likely also causes the repression of transposable elements (Elder et al., 1981; Bilanchone et al., 1993), while defective Nt-acetylation of Orc1, a component of the DNA-binding Origin Recognition Complex (ORC), confers the enhanced expression of sub-telomeric genes (Geissenhöner et al., 2004). The increased expression of mitochondrial genes may represent the cellular response to reduced mitochondrial activity. The identification of the underlying mitochondrial defects and involved NatA substrates requires further investigation.

We speculate that NatA is particularly responsive to environmental cues and confers full acetyltransferase activity only when acetyl-CoA levels are high, i.e. upon growth in glucose rich media. A recent study confirmed changed Nt-acetylation levels of Nat substrates in a protein-specific and not global manner under growth conditions conferring low acetyl-CoA levels (Varland et al., 2018). Overall, the selective effect on fewer protein targets (as compared to NatB) qualifies NatA to play a regulatory role in specific cell responses. As consequence, exponentially growing WT cells may express transposon genes, suppress expression of sub-telomeric genes and allow mating to efficiently protect and reshape the genome, with impact on the genetic diversity of the population, while this is prevented under conditions of perturbed proteostasis and metabolism, with signals transmitted via NatA.

In summary, the two major N-terminal acetyltransferases of yeast, NatA and NatB, have a broad range of functions in maintaining proteome integrity and stress signaling, executed by a clear division of labor between the two enzymes. It is an intriguing possibility that the evolution of NatA and NatB is not only driven by the need for distinct catalytic activities to modify the entire spectrum of N-termini of proteins, but also to elicit enzyme-specific cellular responses.

## Author Contributions

U.A.F., M.Z., G.K., and B.B. conceived and designed the study. U.A.F and M.Z. performed most experiments and analyzed the data. B.H. and L.G. conducted the proteomics experiments and performed data analysis, K.F. performed some ribosome profiling experiments and M.K. and J.B. were involved in the tFT screening. U.A.F., M.Z., G.K., and B.B. wrote the manuscript with input from all authors.

## Acknowledgements

We thank Y. Vainshtein and I. Kats for support with data analysis and B. Zachmann-Brand and D. Ibberson for technical support. Sequencing was carried out at Genomics and Proteomics Core (DKFZ), the Genomics Core (EMBL) and the CellNetWorks Deep Sequencing Core facilities, Heidelberg, Germany. Mass spectrometry was performed at the ETH Zurich and at the ZMBH Heidelberg (Core Facility for Mass Spectrometry & Proteomics). Ludovic Gillet was supported by ETH funding from the group of Ruedi Aebersold. This work was supported by ERC (743118) and DFG (SFB1036) grants to BB.

## Material AND Methods

### Strain and Plasmid Construction

The *Saccharomyces cerevisiae* S288C strain BY4741 served as background for all generated strains in this study. Gene deletion was performed via homologous recombination of the complete indicated *ORF* by one of the selection markers *kanMX4, natMX6, hphNT1, K.l.URA3* as described previously (Janke et al., 2004). Knockouts were confirmed by growth resistance to indicated antibiotics/ auxotrophies and colony PCR.

For complementation with WT or mutant genes, the plasmids pRS416 and p413CYC1 were used. The required genes were generated by PCR using yeast gDNA and respective primer pairs. Double digestion with SpeI-NotI (for pRS416) or SmaI-SalI (for p413CYC1) was performed for the plasmids as well as the PCR products. Corresponding plasmid backbone and insert were purified by agarose gel electrophoresis, ligated with T4 ligase and transformed into chemically competent *E. coli* cells. After selection on ampicillin plates, plasmids were extracted and sequenced to confirm correct gene insertion. Verified plasmids were used for transformation into yeast knockout strains. Mutants of *NAA10* genes were generated by amplification of the generated plasmid using mismatch primers, followed by digestion of the template DNA using Dpn1, transformation, plasmid purification, and sequencing as described earlier.

All yeast strains were grown on full media (YPD) or synthetic defined media (SCD/ SCG) complemented by 2% glucose (SCD) or 2% glycerol (SCG). Heat treated cells were shifted from 30°C to 37°C two doublings prior to harvest. Otherwise cells were grown at 30°C and 130 rpm in liquid culture or at 30°C on agar plates.

Used primers, generated plasmids and strains are given in **Tables S4-S6**, respectively.

### Gene Duplication and Tagging

Genes were duplicated and tagged with a single step PCR protocol following the strategy developed by Huber and coworkers (Huber et al., 2014). Using primers 17-26 (Supplementary Table 5) and plasmid pMaM61 (Khmelinskii et al., 2012), a PCR construct was generated that comprises a tFT-tag, a marker (*natMX4*) to select for successful cloning, and a 3’ and 5’ overhang of 55 nt aligning downstream and upstream to the gene of interest, respectively. This construct was introduced into WT yeast cells by transformation according to standard protocols (Gietz & Schiestl, 2007) and followed by deletion of NatA and NatB, respectively. Verified strains were grown to an OD_600nm_ of 0.5 and harvested by centrifugation at 8,000xg for 1 min at RT. Thorough washing with 1 ml 1x PBS removed residual medium. 100,000 cells of each sample were measured by FACS and analyzed as described previously (Khmelinskii et al., 2016). The median of 2 replicates are shown and statistically analyzed as indicated.

### Growth Assay

The growth behavior of all strains used in this study was analyzed in liquid culture and on solid agar plates. For both methods, a pre-culture was grown to mid-log or early stationary phase and washed three times with dH_2_O (5000xg, 3 min, RT) if medium or carbon source was switched. Spot assays were prepared by stamping 1:10 serial dilutions on respective agar plates.

Main liquid cultures were prepared as 200 μl cultures with a start OD_600nm_ of 0.1 and incubated in clear 12-well or 96-well plates, at 30°C and under continuous shaking (600 rpm). The optical density was measure every 5-20 min in a plate reader (FLUOstar Omega or SPECTROstar Nano, BMG). OD calibration and doubling time calculation was performed as described by Fernandez-Ricaud and coworkers (Fernandez-Ricaud et al., 2007). Each strain was measured in at least three technical and three biological replicates.

### Luciferase Activity Assay

Yeast cells expressing mutant luciferase (Gupta et al., 2011) were grown in SCD medium at 30°C and 130 rpm to mid-log phase. Protein synthesis was stopped by adding cycloheximide to a final concentration of 0.1 mg/ml. Cells were then incubated: 90 min at 37°C as pre-shock, 20 min at 45°C as heat shock and finally at 30°C to allow luciferase refolding. Luciferase activity was measured at different time points: before pre-shock and before heat shock and during refolding phase at 0 min, 15 min, 30 min, 60 min and 120 min. To measure luciferase activity, 100 μl of cell culture was mixed with 100 μl of 125 μM luciferin and luciferase activity was measured for 10 s (Lumat LB 9507, Berthold Technologies GmbH & Co. KG). Measurements were normalized to the optical density (OD_600nm_). The experiment was done in replicates.

### ^35^S-Methionine Incorporation

The experiment was performed as described previously (Nillegoda et al., 2010), with a few modifications. In brief, yeast cells were grown in SCD to mid-log phase, washed twice in dH_2_O, and resuspended in synthetic dropout medium lacking methionine to a final concentration of 6 OD_600nm_ / ml. ^35^S-methionine was added to a final concentration of 100 μCi/ml. Pulse-labeling was conducted for 10 min at 30°C while shaking. Incorporation of ^35^S-methionine was measured by mixing 400 μl labeled culture 1:1 (v/v) with ice-cold 20% TCA. Cells were pelleted, washed twice in ice-cold acetone then resuspended in 200 μl ice-cold extraction buffer (50 mM Tris-HCl pH 7.5, 1 mM EDTA, 1x protease inhibitors (Complete EDTA-free, Roche), 1% SDS). Equal volume of glass beads (500 μm diameter) was added to the cells, followed by vortexing for 40 s at 6.0 m/s (FastPrep 24, MP Biomedical). The extracts were quantified for ^35^S incorporation using a scintillation counter (1900 TR β, Packard).

### Mitochondrial Staining using MitoTracker

Mitochondria were stained and quantified using the dye MitoTracker Orange CMTMRos (Invitrogen). Yeast cells were grown in SCD medium, at 30°C to OD_600nm_ = 0.5, harvested by centrifugation at 1,150xg for 3 min and resuspended in 750 μl 250 nM MitoTracker dissolved in SCD. A negative control without MitoTracker in the growth medium was performed in parallel. The subsequent incubation at 30°C and 150 rpm was performed for 15 min and as all following incubation steps under light-protected conditions. Cells were washed twice with 500 μl 1x PBS and incubated for 10 min in 4% paraformaldehyde in the growth medium. Four times washing in 500 μl 1x PBS (1,150xg, 3 min) prepared the fixed cells for quantitative analysis by FACS (BD FACSCanto, BD Biosciences). The fluorescent MitoTracker dye was detected using the mCherry channel (excitation: 587 nm, emission: 610 nm) and the cell size by side scatter.

### mRNA Sequencing and Ribosome Profiling

100 OD_600nm_ of mid-log culture was filtered (All-Glass Filter 90mm, Millipore), flash frozen in liquid nitrogen and lysed by mixer milling (2 min, 30 Hz, MM400, Retsch) with 600 μl lysis buffer (20 mM Tris-HCl pH 8.0, 140 mM KCl, 6 mM MgCl_2_, 0.1% NP-40, 0.1 mg/ml cycloheximide, 1 mM PMSF, 2x protease inhibitor cocktail (Complete EDTA-free, Roche), 0.02 U/ml DNaseI (recombinant DNaseI, Roche), 20 mg/ml leupeptin, 20 mg/ml aprotinin, 10 mg/ml E-64, 40 mg/ml bestatin). Thawed cell lysates were cleared from cell debris by centrifugation (20,000xg, 5 min, 4°C). From this sample, 200 μl total RNA were taken for RNA sequencing and 1000 μl total RNA were used for further nuclease digestion and ribosome profiling. Ribosome footprints were generated by a digest with RNase I (Ambion, 1 hr, 25°C, 650 rpm) which was stopped by 10 μl SUPERase-In (Ambion). Monosome fractions were generated by polysome profiling (see below) and pooled for each sample. RNA from both RNAseq and RP samples was extracted by hot phenol extraction as described previously (Döring et al., 2017). 10 μg of extracted RNA of the RNAseq sample was in addition depleted for rRNA using the kit RiboMinus, Yeast Module (Invitrogen) and fragmented for 30 min using the NEBNext Magnesium RNA Fragmentation Module (NEB). Deep sequencing libraries for RNAseq and RP samples were prepared exactly following previously published protocols (Döring et al., 2017; Shiber et al., 2018). The generated libraries were sequenced on a HiSeq 2000 system (Illumina).

All deep sequencing data were trimmed, aligned to non-coding elements (excluded) and to the yeast genome R64.2.1 (included). The used parameters for the publically available tools Trim Galore, Bowtie and Tophat as well as the self-written scripts for the subsequent analyses were published previously (Galmozzi et al., 2019). In short, RNAseq reads and ribosome footprints were allowed to align with a maximum of 2 mismatches and must have a length between 23 and 41 nt. The approximate center of the ribosome was statistically estimated for the ribosome footprints by (virtually) cutting 11 nt of either side of the footprint (‘center weighting’) while RNAseq reads were evenly distributed over the whole length. Genes with ≥64 raw reads in at least one of the two replicates were included in subsequent analyses. Differentially expressed genes were altered more than 2-fold in both replicates. GO term analyses were performed with Perseus version 1.4.1.3 (Tyanova et al., 2016), using an false discovery rate (FDR) of 2%.

### Polysome Profiling

Cells were grown, harvested and lysed as for mRNA sequencing and ribosome profiling. Thawed cell lysates were cleared by centrifugation (20,000xg, 5 min, 4°C) and 500 μg total RNA was loaded onto 10-50% sucrose gradients. Monosome vs. polysome separation was performed by ultracentrifugation for 2.5 hrs at 35,000 rpm and 4°C (SW40-rotor, Sorvall Discovery 100SE Ultracentrifuge). The centrifuged gradients were fractionated by a piston gradient fractionator (BioComp) and absorbance at 254 nm was analyzed.

### Stable Isotope Labeling with Amino Acids in Cell Culture (SILAC)

Quantitative proteomics of total lysates or protein aggregates by SILAC-based mass spectrometry were performed with strains lacking the genes *ARG4* and LYS1 and the use of heavy (^13^C_6_, ^15^N_2_) or light (^12^C_6_, ^14^N_2_) isotopes L-arginine and L-lysine in SCD media. To avoid any potential label-dependent effects, replicates of mutant and WT strains were inversely cultivated in media with heavy or light isotopes and labeling efficiency was separately tested and exceeds 95% for all strains and growth conditions.

At least 50 OD_600nm_ of culture in mid-log phase was harvested by centrifugation at 3,000xg and 4°C for 3 min. The cell pellet was resuspended with 600 μl lysis buffer (20 mM Tris-HCl pH 8.0, 140 mM KCl, 6 mM MgCl_2_, 0.1% NP-40, 0.1 mg/ml cycloheximide, 1 mM PMSF, 1x protease inhibitor cocktail (Complete EDTA-free, Roche), 0.02 U/ml DNaseI (recombinant DNaseI, Roche), 20 mg/ml leupeptin, 20 mg/ml aprotinin, 10 mg/ml E-64, 40 mg/ml bestatin) and frozen as droplets in liquid nitrogen. Corresponding heavy and light labeled samples i.e. WT and Nat mutant strains were mixed 1:1 and lysed in frozen state by mixer milling with 30 Hz for 2 min (MM400, Retsch). Centrifugation at 300xg for 20 min (4°C) removed cell debris and proteins of the supernatant were precipitated using 12% trichloroacetic acid. The protein pellet was resuspended in SDS sample buffer and run on a 12% SDS gel. Gel pieces were reduced with DTT, alkylated with iodoacetamid and digested with trypsin using the DigestPro MS platform (Intavis AG) following the protocol described by Shevchenko and colleagues (Shevchenko et al., 2006).

Peptides have then been analyzed by liquid chromatography-mass spectrometry (LCMS) using an UltiMate 3000 LC (Thermo Scientific) coupled to either an Orbitrap Elite or a Q-Exactive mass spectrometer (Thermo Scientific). Peptides analyzed by the Orbitrap Elite have been loaded on a C18 Acclaim PepMap100 trap-column (Thermo Fisher Scientific) with a flow rate of 30 μl/min 0.1% TFA. Peptides were eluted and separated on an C18 Acclaim PepMap RSLC analytical column (75 μm x 250 mm) with a flow rate of 300 nl/min in a 2 hrs gradient of 3% buffer A (0.1% formic acid, 1% acetonitril) to 40% buffer B (0.1% formic acid, 90% acetonitrile). MS data were acquired with an automatic switch between a full scan and up to 30 data-dependent MS/MS scans. Peptides analyzed on the Q-Exactive have been directly injected to an analytical column (75 μm x 300 mm), which was self-packed with 3 μm Reprosil Pur-AQ C18 material (Dr. Maisch HPLC GmbH) and separated using the same gradient as described before. MS data were acquired with an automatic switch between a full scan and up to 15 data-dependent MS/MS scans.

Data analysis was carried out with MaxQuant version 1.5.2.8 (Cox & Mann, 2008) using standard settings for each instrument type and searched against a yeast specific database extracted from UniProt (UniProt Consortium). Carbamidomethylation of cysteine was specified as fixed modification; Oxidation of methionine and acetylation of protein N-termini was set as variable modification. ‘Requantify’ as well as ‘Match Between Runs’ options were both enabled.

Results were filtered for an 1% FDR on peptide spectrum match (PSM) and protein level. MaxQuant output files have been further processed and filtered using self-compiled R-scripts.

### Isolation of Protein Aggregates

Aggregated proteins were isolated following previously published protocols (Koplin et al., 2010). In summary, the cell pellet of 50 OD_600nm_ of culture in mid-log phase was resuspended in 100 μl lysis buffer (20 mM Na-phosphate pH 6.8, 10 mM DTT, 1 mM EDTA, 0.1% Tween, 1 mM PMSF, 1x protease inhibitor cocktail (Complete EDTA-free, Roche), 3 mg/ml zymolyase T20) and incubated for 20 min at 25°C and 750 rpm. The resuspended cells were kept on ice, followed by tip sonication (Branson, 8x at level 4 and duty cycle 50%) and centrifugation for 20 min at 200xg and 4°C. The supernatant was transferred to a new tube. All supernatants were adjusted to identical protein concentrations using Bradford. Aggregated proteins were pelleted for 20 min at 16,000xg, 4°C. The supernatants were removed and aggregated proteins were washed twice with 600 μl wash buffer (20 mM Na-phosphate pH 6.8, 1 mM PMSF, 1x protease inhibitor cocktail (Complete EDTA-free, Roche), 2% NP-40) by sonication (Branson, 6x at level 4 and duty cycle 50%) and collected by centrifugation for 20 min at 16,000xg, 4°C. A further washing step was performed with wash buffer without NP-40 and 4x sonication at level 2 and duty cycle 65%. The samples were boiled in SDS sample buffer and separated on a 14% SDS gel. For subsequent identification and quantification by SILAC mass spectrometry, aggregates from WT and mutant strains were mixed 1:1 in respect to the initial total lysate protein concentration prior to SDS-PAGE. Proteins with more than two-fold elevated levels in the aggregates isolated from the mutants relative to WT were counted as aggregated protein in the respective mutant, and were exclusively included in the downstream analysis. For detection of newly synthesized aggregated protein, cells were grown in synthetic defined medium, and shifted to synthetic defined medium lacking methionine. ^35^S-methionine was added to a final concentration of 100 μCi/ml. Cells were pulse labeled for 5 min, and chased with 20 mM cold methionine. Protein aggregates were isolated as described above.

### SWATH-based quantitative proteomics

Yeast cells were grown in triplicates to mid-log phase, quenched by addition of trichloroacetic acid (TCA) to a final concentration of 6.25% (v/v) and incubated on ice for 10 min. The culture was harvested by centrifugation for 5 min at 1,500xg and 4°C. The cell pellet was washed twice with 10 ml ice-cold acetone. The cell lysis was performed by bead-beating in 8 M urea, 100 mM NH_4_HCO_3_, 5 mM EDTA at pH 8.0 and the extracted proteins were reduced, alkylated and digested with trypsin as described previously (Selevsek et al., 2015). The peptides were desalted with C18 cartridges and injected using an Eksigent nanoLC Ultra on a 20cm-long new Objective emitter packed with Magic C18 AQ 3 μm resin (Selevsek et al., 2015). The MS data was acquired by a Sciex 5600 TripleTOF operating either in DDA mode for library generation or in SWATH mode for quantitation. The DDA spectral library was generated as described in Schubert et al., 2015. The SWATH MS data was extracted with openSWATH (Röst et al., 2014) and aligned with TRIC (Röst et al., 2016). The final protein matrix and statistical analysis was generated using mapDIA (Teo et al., 2015). The raw data and analysis results have been deposited to the ProteomeXchange Consortium via PRIDE (Perez-Riverol et al., 2019) with the dataset identifier PXD015217.

### Tandem Fluorescent Timer Assay

Tandem fluorescent timer analysis was performed as described (Khmelinskii et al., 2014), with the following specifications: The screen was done in 1536 format using pinning robotics (Singer instruments) with three biological replicates per clone. Haploids that have both tFT gene fusion and *natB*Δ mutation were grown on synthetic complete media without histidine (for haploid selection) for 1 day followed by high throughput fluorescence measurements of colonies for sfGFP and mCherry intensities. To test the effect of heat stress, plates were incubated at 30°C for 1 day then shifted to 37°C for 1 day followed by high throughput fluorescence measurement of colonies using Infinite M1000 or Infinite M1000 Pro plate readers equipped with stackers for automated plate loading (Tecan) and custom temperature control chambers. Data analysis for the measured fluorescence intensities was performed as described (Khmelinskii et al., 2014).

