## Supplementary Figures S1-S11 for "N^α^-terminal acetylation of proteins by NatA and NatB serves distinct physiological roles in *Saccharomyces cerevisiae*"

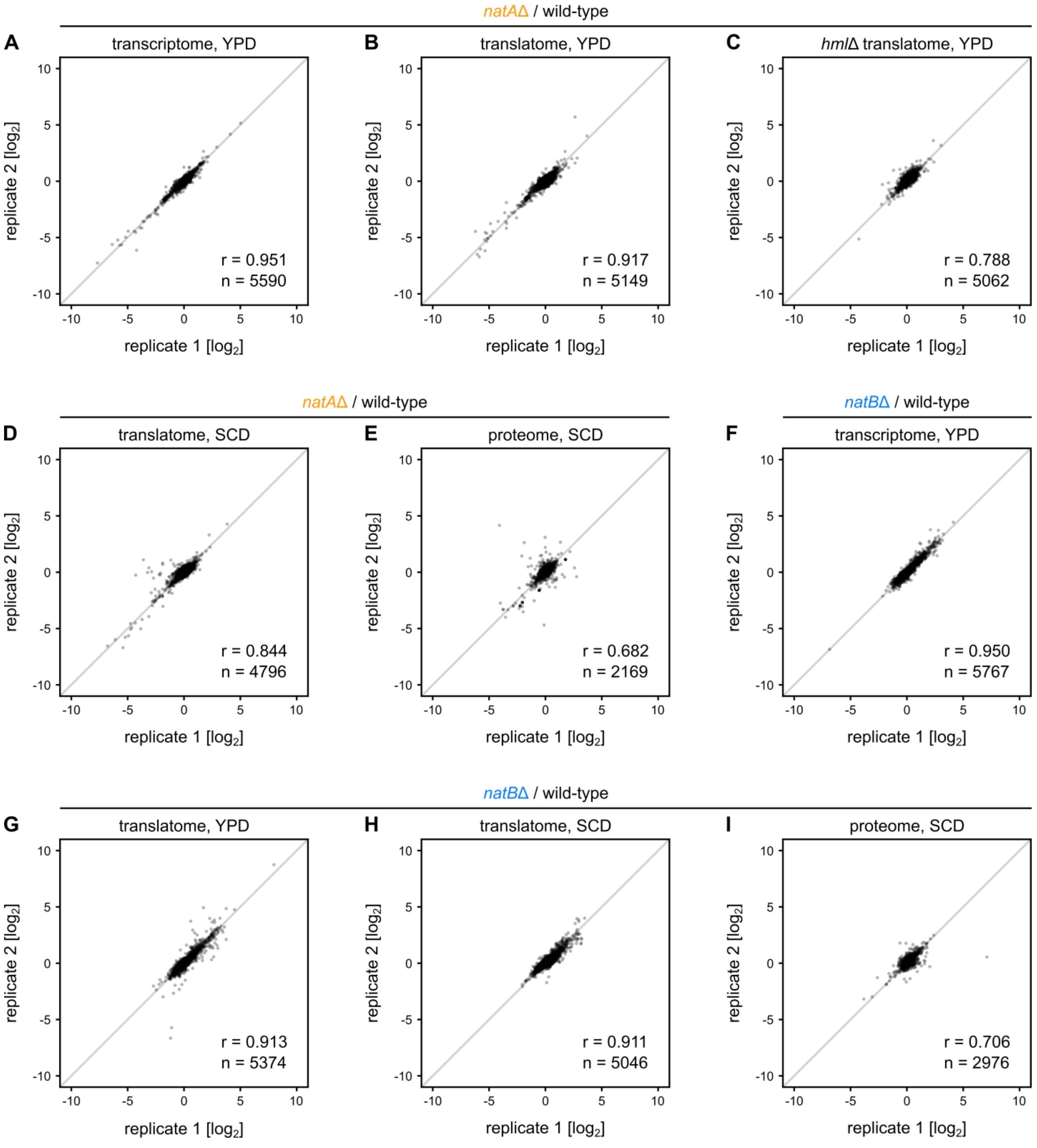


**Figure S1. Reproducibility of RNAseq, ribosome profiling and SILAC-based proteomics**
Relative changes of mRNA levels (A, F), protein synthesis rates (B, C, D, G, H) or protein levels (E, I) between Nat deletion mutants and WT cells. Pearson correlation coefficient (r) is calculated for the given number (n) of quantified transcripts or proteins.

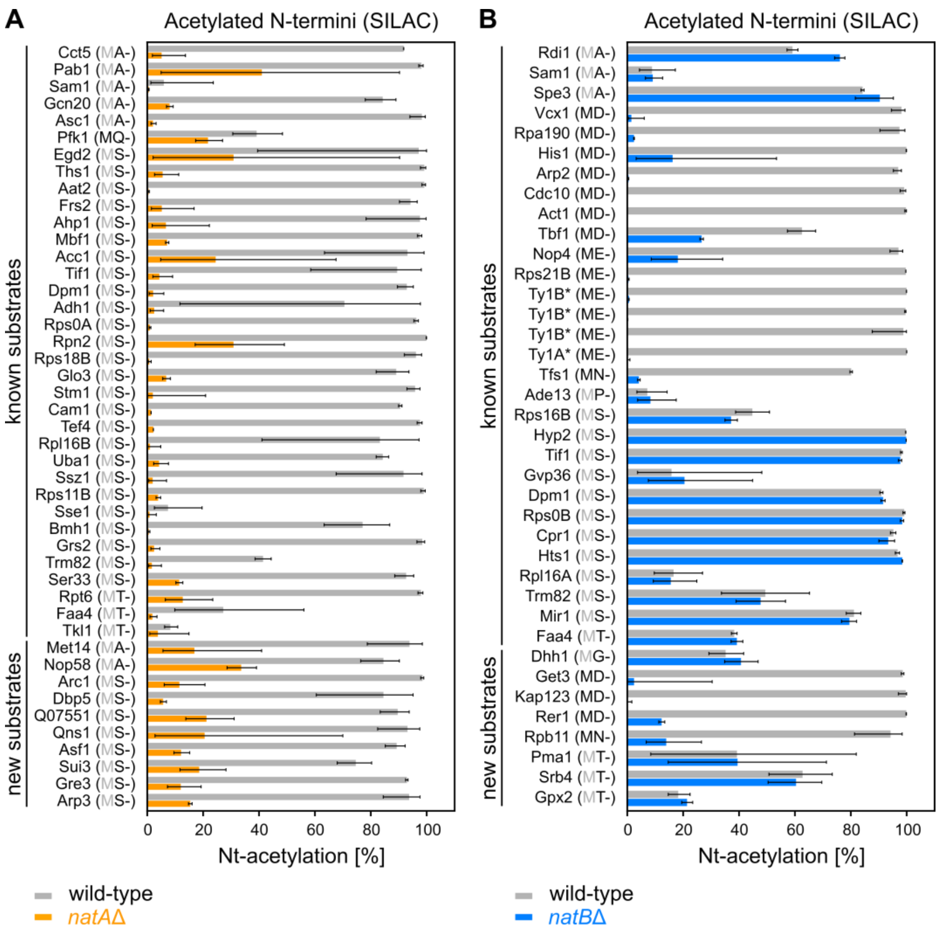


**Figure S2. Loss of Nt-acetylation of specific proteins in *natA*Δ and *natB*Δ mutant cells**

Percentages of Nt-acetylation in WT vs. *natA*Δ (A) or *natB*Δ (B) mutant cells for proteins where both N-terminally acetylated and non N-terminally acetylated forms of the same peptides. The sequence of the first two N-terminal amino-acids is listed in brackets.

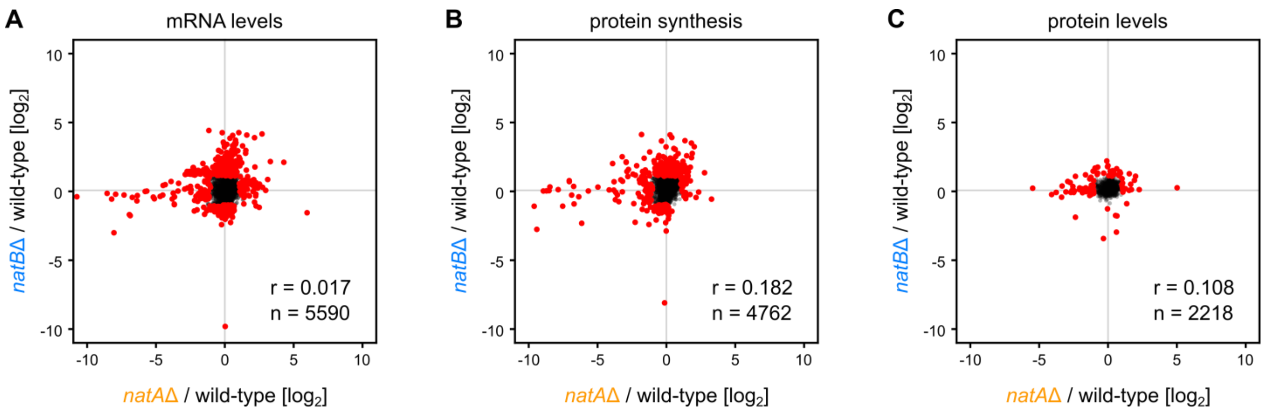


**Figure S3. Correlation of NatA- and NatB-induced changes**
Knockout-induced changes at the level of mRNA (A), protein synthesis (B) or protein (C) compared between NatA and NatB. Pearson correlation coefficient (r) is computed for all quantified transcripts or proteins (n). Red dots indicate transcripts or proteins that are more than 2-fold altered in at least one of the two data sets.

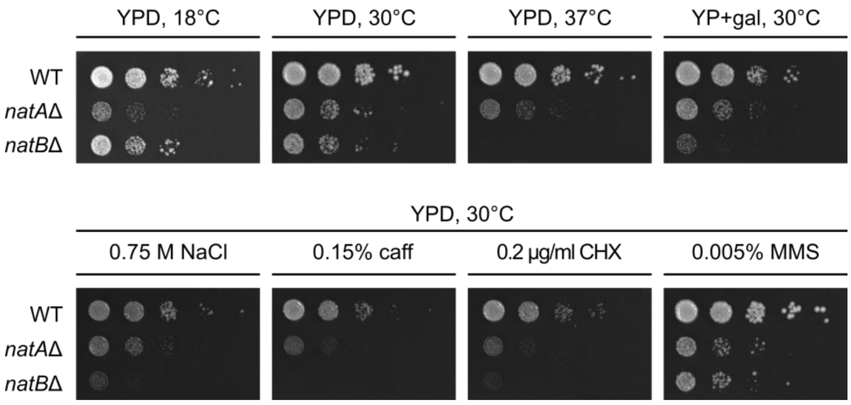


**Figure S4. Growth phenotype of NatA and NatB knockouts on solid agar**
Growth of WT and Nat mutant strains on solid agar plates and different growth conditions. Cells are serially diluted 1:10 with an initial $OD_{600nm}$ of 1.0, spotted on plates with the indicated carbon source or supplement. Plates are incubated at the given temperature for 2 days or at 18°C for 5 days. gal – 2% galactose, caff – caffeine, CHX – cycloheximide, MMS – methyl methanesulfonate.

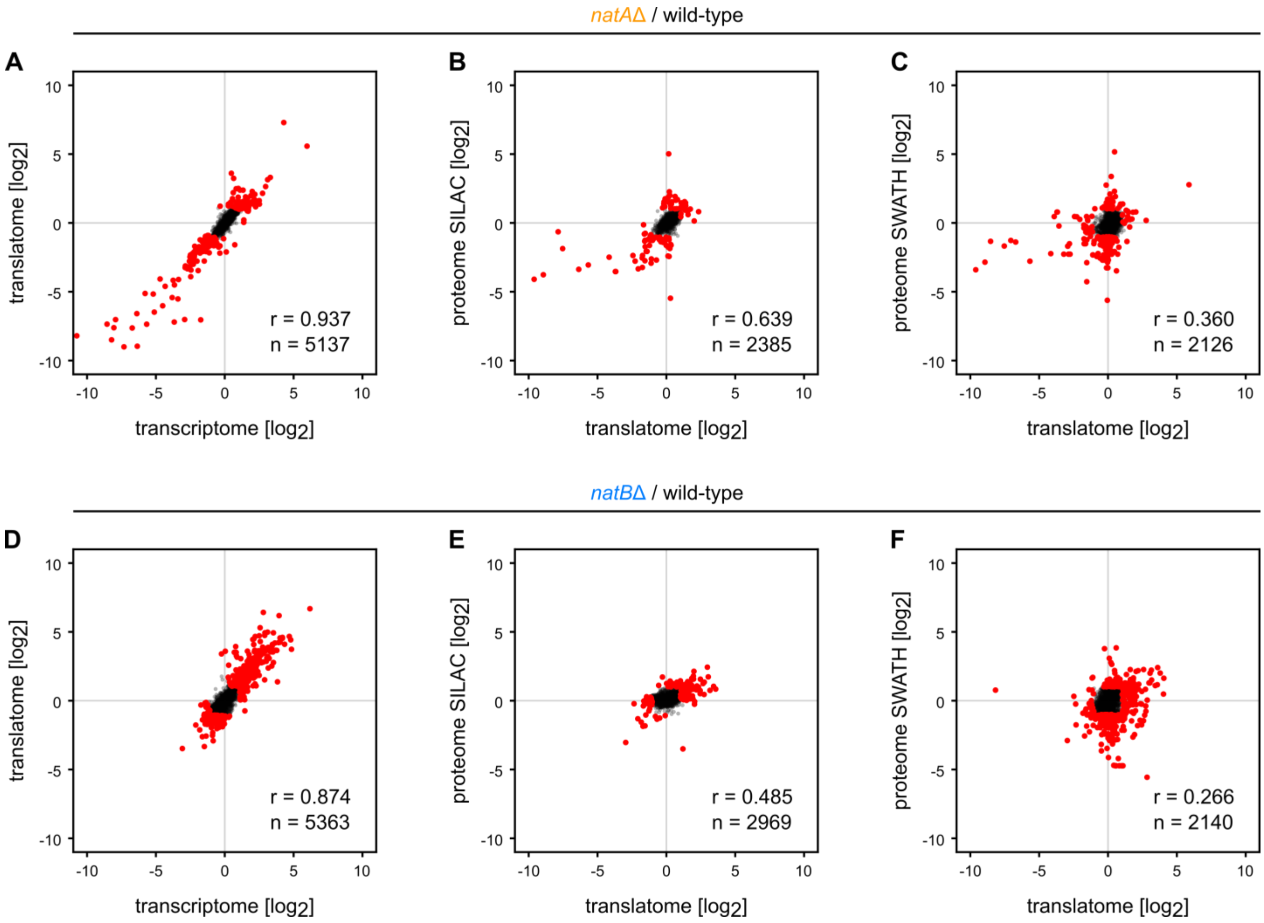


**Figure S5. Correlation between high-throughput omics data sets for *natA*Δ and *natB*Δ mutant cells** Pairwise comparison of deletion induced changes in mRNA levels and protein synthesis (A, D) and protein synthesis and levels (B, C, E, F) between WT and *natA*Δ and *natB*Δ mutant cells, respectively. Pearson correlation coefficient (r) is computed for the $log_2$-transform of all quantified transcripts or proteins present in both data sets (n). Red dots indicate transcripts or proteins more than 2-fold altered in at least one of the two data sets.

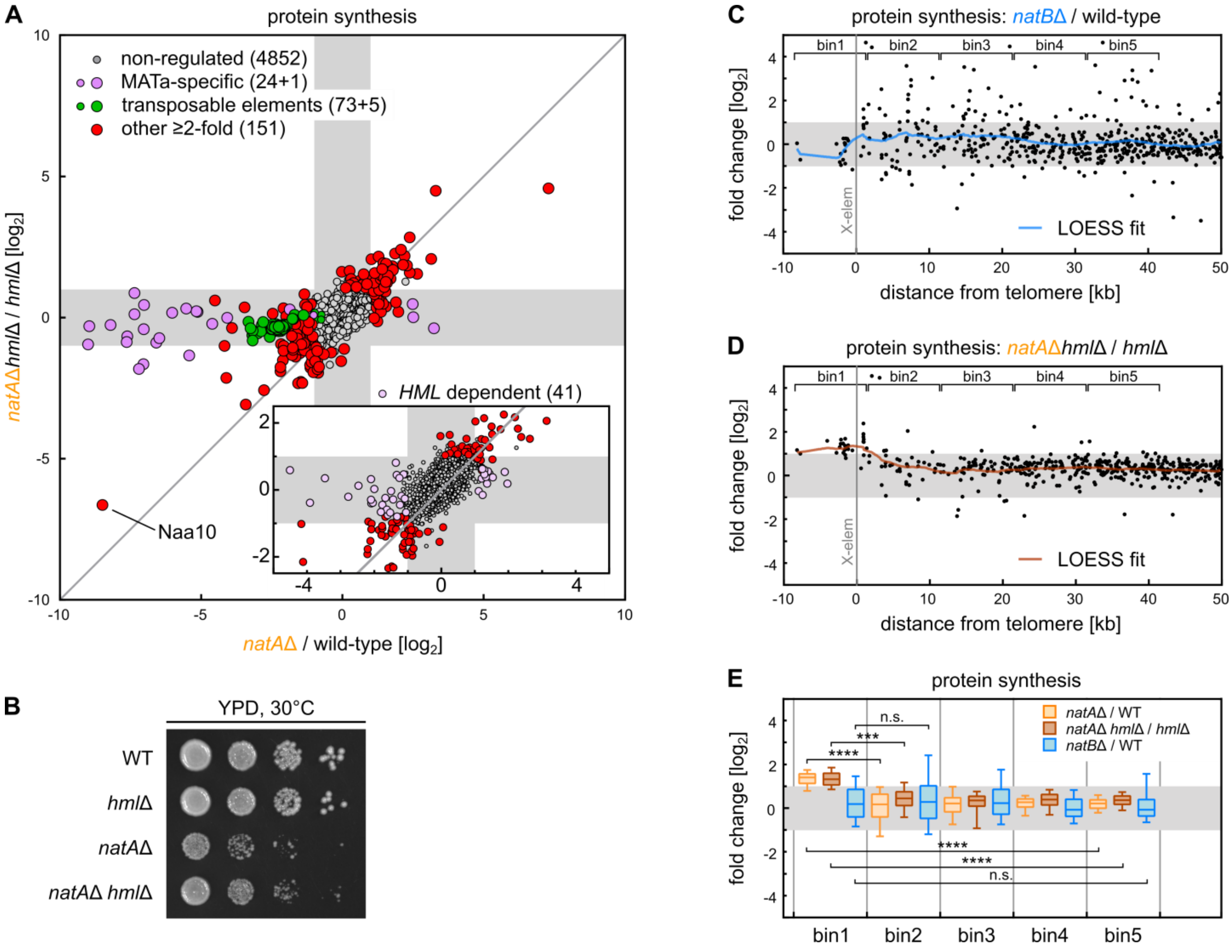


**Figure S6. Characterization of *HML*-mediated protein synthesis regulation in *natA*Δ cells and influence of gene localization in proximity to telomeres**

(A) Scatter plot as in Figure 2A. The insert shows the same data without MATa-specific proteins and transposable elements, highlighting 41 proteins regulated in *natA*Δ in an *HML*-dependent way. (B) Serially diluted (1:10) *natA*Δ and *hml*Δ mutant strains with an initial $OD_{600nm}$ of 1.0, spotted on YPD plates and incubated for 2 days at 30°C. (C, D) As in Figure 2D, but for *natB*Δ mutant or *natA*Δ *hml*Δ double knockout. (E) Data from Figures 2D, S2C, and S2D binned (10 kb) and statistically compared between bins using the two-sided Student's t-test.

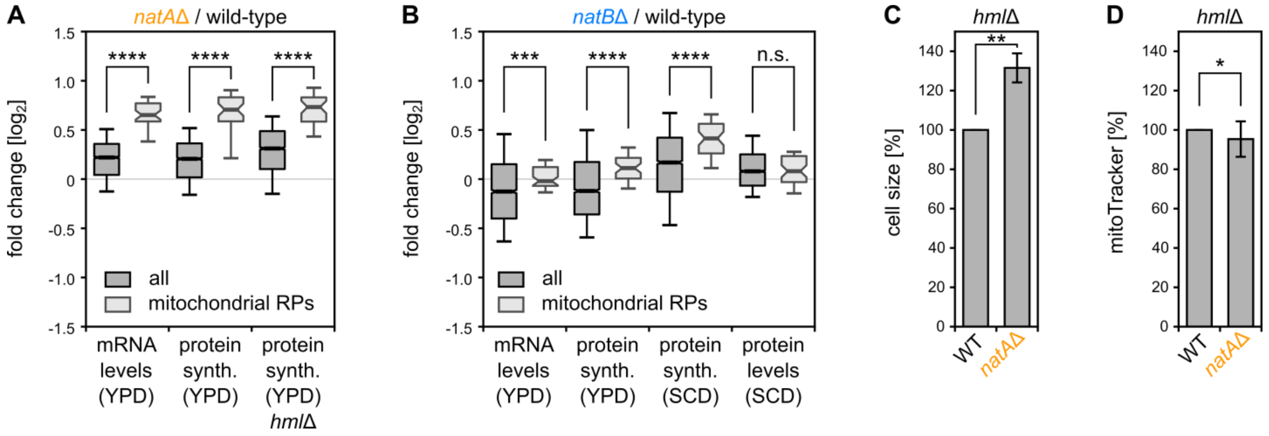


**Figure S7. Mitochondrial analysis of Nat mutants and influence of the *HML* cassette**

mRNA levels, synthesis and steady state levels of proteins in *natA*Δ and *natA*Δ *hml*Δ (A) or *natB*Δ (B) mutant cells compared to WT grown in full (YPD) or synthetically defined media (SCD) determined by RNAseq, RP and SILAC, respectively. $Log_2$-transforms of all proteins (2372 and 2969, respectively) and mitochondrial ribosomal proteins (38 and 47, respectively) are statistically compared by Mann-Whitney U test. (C, D) FACS analysis as in Figures 3B and 3C in the *hml*Δ background.

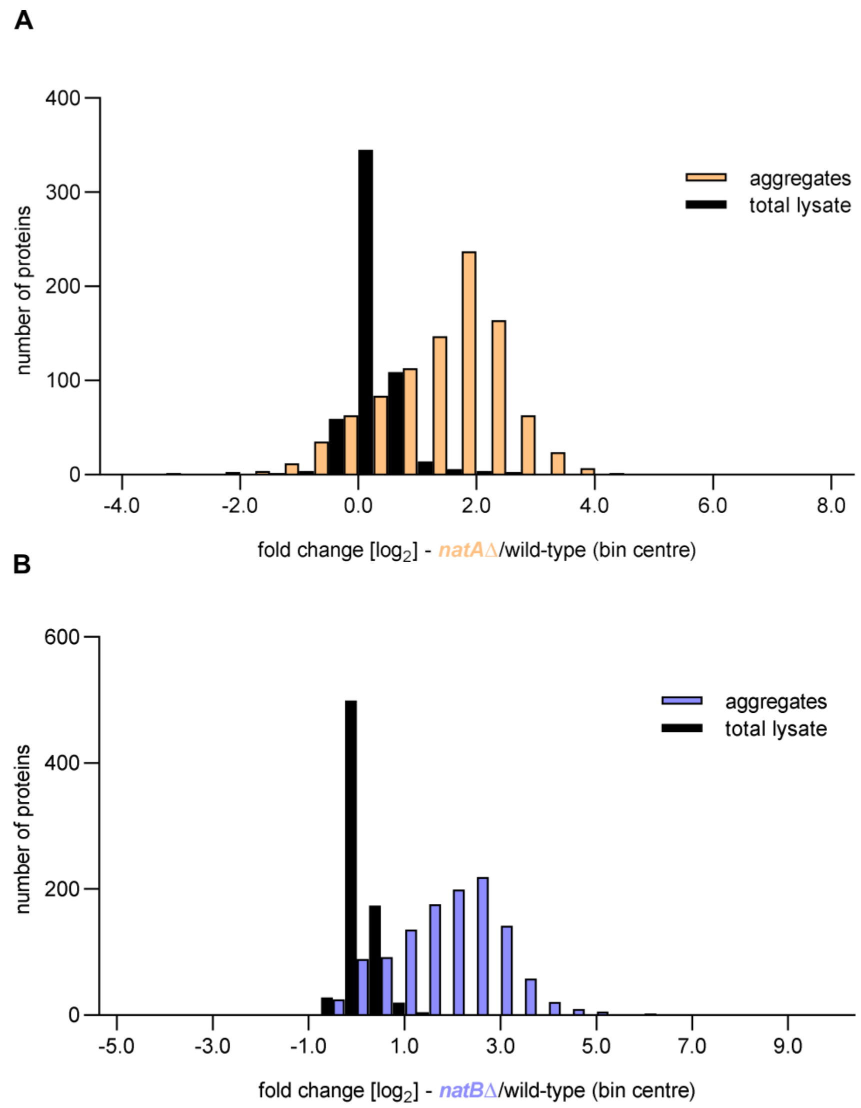


**Figure S8. Global protein aggregation is not driven by altered of total protein levels in the mutants relative to WT**

Comparison of the distribution of protein level changes in *natA*Δ (A) and *natB*Δ (B) mutant cells relative to WT, in total lysate and aggregates.

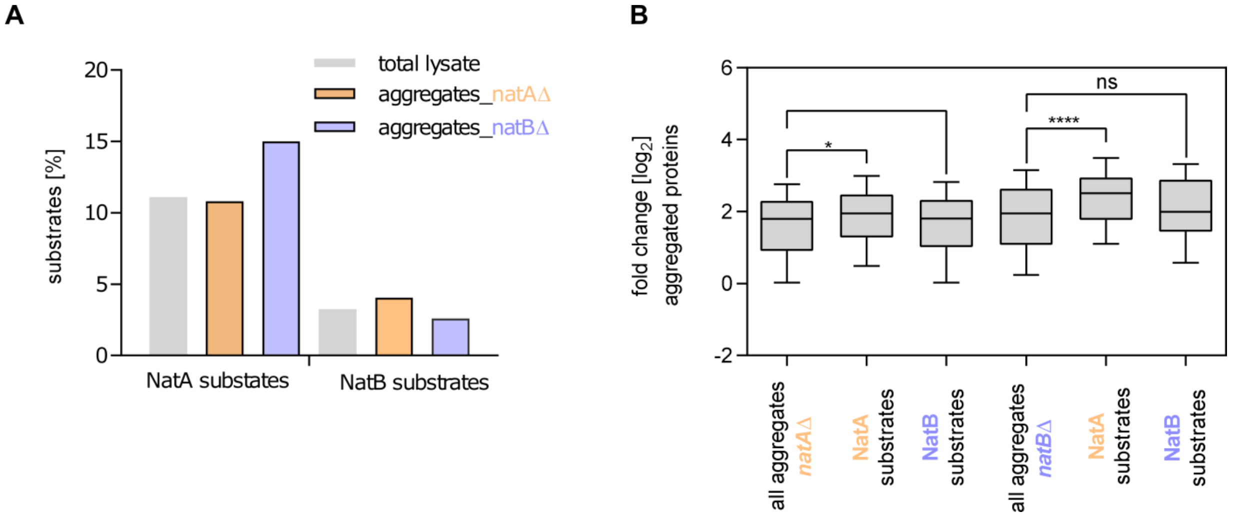


**Figure S9. Global protein aggregation is not specific to Nat substrates**
(A) The percentage of NatA and NatB substrates among quantified proteins in the total lysate or aggregated proteins in *natA*Δ and *natB*Δ mutant compared to WT cells. (B) Comparison of the distribution of protein level changes for verified NatA and NatB substrates, versus all aggregated proteins in *natA*Δ and *natB*Δ mutant cells relative to WT, for the aggregate fraction.

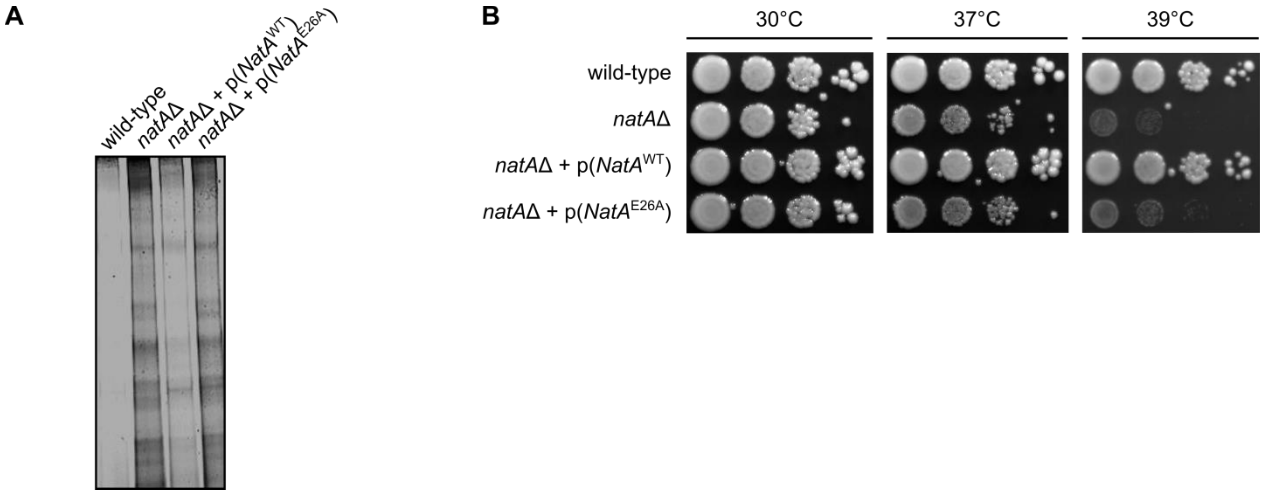


**Figure S10. Lack of proteins Nt- acetylation activity causes global protein aggregation**
(A) Comparison of levels of isolated protein aggregates from WT, *natA*Δ, and *natA*Δ expressing plasmid-encoded WT NatA catalytic subunit (denoted $NatA^{WT}$), or the catalytically dead mutant of the NatA catalytic subunit (denoted $NatA^{E26A}$), visualized by SDS-PAGE and Coomassie staining. (B) Growth of WT, *natA*Δ, and *natA*Δ expressing plasmid-encoded WT NatA catalytic subunit (denoted $NatA^{WT}$), or the catalytically dead mutant of the NatA catalytic subunit (denoted $NatA^{E26A}$) on YPD agar plates.

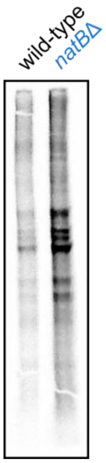


**Figure S11. Aggregation of newly synthesized proteins upon loss of Nt-acetylation**
$^{35}$S-methionine labeled protein aggregates isolated from *natB*Δ mutant cells and WT after 5 min pulse labeling analyzed by SDS-PAGE and autoradiography.
